# Fetal liver macrophages contribute to the hematopoietic stem cell niche by controlling granulopoiesis

**DOI:** 10.1101/2023.02.21.529403

**Authors:** Amir Hossein Kayvanjoo, Iva Splichalova, David Alejandro Bejarano, Hao Huang, Katharina Mauel, Nikola Makdissi, David Heider, Nora Reka Balzer, Collins Osei-Sarpong, Kevin Baßler, Joachim L. Schultze, Stefan Uderhardt, Eva Kiermaier, Marc Beyer, Andreas Schlitzer, Elvira Mass

**Affiliations:** Developmental Biology of the Immune System, Life & Medical Sciences (LIMES) Institute, University of Bonn; 53115 Bonn, Germany; Quantitative Systems Biology, Life & Medical Sciences (LIMES) Institute, University of Bonn; 53115 Bonn, Germany; Immunogenomics & Neurodegeneration, Deutsches Zentrum für Neurodegenerative Erkrankungen (DZNE) e.V., Bonn, Germany; Genomics & Immunoregulation, LIMES Institute, University of Bonn, Bonn, Germany; Systems Medicine, Deutsches Zentrum für Neurodegenerative Erkrankungen (DZNE) e.V., Bonn, Germany; PRECISE Platform for Single Cell Genomics and Epigenomics, DZNE and University of Bonn, Bonn, Germany; Deutsches Zentrum für Immuntherapie (DZI), Friedrich Alexander University Erlangen-Nuremberg and Universitätsklinikum Erlangen, Erlangen, Germany; Department of Internal Medicine 3-Rheumatology and Immunology, Friedrich Alexander University Erlangen-Nuremberg and Universitätsklinikum Erlangen, Erlangen, Germany; Immune and Tumor Biology, Life & Medical Sciences (LIMES) Institute, University of Bonn; 53115 Bonn, Germany

## Abstract

During embryogenesis, the fetal liver becomes the main hematopoietic organ, where stem and progenitor cells as well as immature and mature immune cells form an intricate cellular network. Hematopoietic stem cells (HSCs) reside in a specialized niche, which is essential for their proliferation and differentiation. However, the cellular and molecular determinants contributing to this fetal HSC niche remain largely unknown. Macrophages are the first differentiated hematopoietic cells found in the developing liver, where they are important for fetal erythropoiesis by promoting erythrocyte maturation and phagocytosing expelled nuclei. Yet, whether macrophages play a role in fetal hematopoiesis beyond serving as a niche for maturing erythroblasts remains elusive. Here, we investigate the heterogeneity of macrophage populations in the fetal liver to define their specific roles during hematopoiesis. Using a single-cell omics approach combined with spatial proteomics and genetic fate-mapping models, we found that fetal liver macrophages cluster into distinct yolk sac-derived subpopulations and that long-term HSCs are interacting preferentially with one of the macrophage subpopulations. Fetal livers lacking macrophages show a delay in erythropoiesis and have an increased number of granulocytes, which can be attributed to transcriptional reprogramming and altered differentiation potential of long-term HSCs. Together, our data provide a detailed map of fetal liver macrophage subpopulations and implicate macrophages as part of the fetal HSC niche.

## Introduction

Macrophages are found in all adult organs, where they perform essential functions during inflammatory responses as well as in tissue homeostasis, such as tissue remodelling, phagocytosis of apoptotic cells, and production of cytokines and growth factors. Work in mice showed that most macrophages originate from erythro-myeloid progenitors (EMPs) in the yolk sac and are long-lived, and that their maintenance in many adult tissues does not rely on definitive hematopoiesis (Gomez Perdiguero et al., 2015; Hoeffel et al., 2015; Mass, 2018). All developing tissues are initially colonized by circulating pre-macrophages (pMacs), which immediately differentiate into tissue-specific macrophages (Mass et al., 2016; Stremmel et al., 2018), therefore, macrophages are an integral part of organogenesis.

In adult tissues, macrophages inhabit distinct anatomical niches within organs, e.g., the lung harbours alveolar and interstitial macrophage populations (Aegerter et al., 2022), while macrophages in the adult liver are divided into Kupffer cells, liver capsular, central vein and lipid-associated macrophages (Guilliams and Scott, 2022). Within these niches, resident macrophages adapt to their tissue environment and perform specific tasks to maintain organ function, such as mucus clearance by alveolar macrophages or phagocytosis of red blood cells by Kupffer cells. However, the role of macrophages in organ development and function as well as their heterogeneity during embryogenesis is less well understood.

One of the few well-known functions of fetal macrophages is their involvement in erythroblast maturation. In the fetal liver, erythroblastic island (EI) macrophages serve as niches for erythroblasts. During erythropoiesis, primitive and definitive erythroblasts directly interact with EI macrophages, where they undergo final steps of maturation, including expelling of their nucleus, which is phagocytosed by EI macrophages (Palis, 2017, 2014). Previous studies identified EI macrophage heterogeneity in the fetal liver at embryonic day (E)13.5/E14.5 (Li et al., 2019; Mukherjee et al., 2021; Seu et al., 2017). EI macrophages were shown to express different levels of cell adhesion proteins such as Vcam1, CD169, and CD163, as well as other proteins that are important for their function as EI niche (e.g., Epor, Klf1, EMP, DNaseII), thereby promoting erythropoiesis (Li et al., 2019; Mariani et al., 2019; May and Forrester, 2020; Mukherjee et al., 2021). In addition to erythropoiesis, the E13.5/E14.5 fetal liver is at the peak of hematopoiesis, providing a niche for the expansion and differentiation of other hematopoietic stem and progenitor cells (Lewis et al., 2021). Thus, the liver is a complex cellular interaction network where the role of macrophages in other hematopoietic developmental processes, such as myelopoiesis, is not fully understood.

Recent evidence suggests that one of the core macrophage functions is the support of stem and progenitor cell functionality in different tissues. For instance, pericryptal macrophages in the gut interact with epithelial progenitors, thereby promoting their proliferation and differentiation via Wnt, gp130, TLR4, or NOX1 signalling (Delfini et al., 2022). In the bone marrow, distinct macrophage subpopulations contribute directly and indirectly to the hematopoietic stem cell (HSC) niche. Osteoclasts are a highly specialized EMP-derived macrophage population responsible for bone resorption, whose function is necessary to generate the bone marrow and, thereby, the postnatal HSC niche (Jacome-Galarza et al., 2019). Co-culture experiments using different combinations of hematopoietic cells indicate that osteomacs support the HSC niche in synergy with megakaryocytes (Mohamad et al., 2017). DARC^+^ macrophages are in direct contact with CD82^+^ long-term (LT)-HSCs in the endosteal and arteriolar niches, where they contribute to LT-HSC dormancy via maintenance of CD82 expression (Hur et al., 2016). Additional macrophage depletion studies via clodronate and diphtheria toxin indicate that CD169^+^ macrophages in the bone marrow promote HSC retention by acting specifically on the Nestin^+^ HSC niche (Chow et al., 2011), as well as steady-state and stress erythropoiesis (Chow et al., 2013).

During embryogenesis, macrophages have been shown to be important for the development and proliferation of hematopoietic stem- and progenitor cells in mice. In the mouse, the first HSCs develop from the aorta-gonad-mesonephros (AGM) region starting at E10.5. Here, a CD206^+^ macrophage population actively interacts with nascent and emerging intra-aortic HSCs (Mariani et al., 2019). Macrophage depletion studies via clodronate liposomes and the Csf1r inhibitor BLZ945 using AGM explants indicate that the presence of macrophages in the AGM is influencing HSC production (Mariani et al., 2019). Furthermore, co-culture studies of a mixture of hematopoietic stem- and multipotent-progenitor cells (HSC/MPP) with macrophages isolated from the fetal liver as well as clodronate depletion studies during embryogenesis suggest that fetal liver macrophages promote HSC/MPP proliferation (Gao et al., 2021). Indeed, immunofluorescent stainings of F4/80^+^ macrophages and CD150^+^ cells (Gao et al., 2021) indicate that macrophages could also be part of the LT-HSC niche in the fetal liver, thereby influencing the proliferation and differentiation of stem cells at the top of the HSC hierarchy.

Studying stem cell niches and defining the molecular factors that promote stem cell proliferation and differentiation is essential to build a robust and reproducible *in vitro* system that may serve as a universal source of functional stem cells and/or immune cells for therapeutic purposes. This has already been achieved for induced pluripotent stem cells (iPSCs), which can be infinitely expanded and differentiated into the cell type of interest, thereby representing a safe product to treat diseases (Yamanaka, 2020). In contrast, a universal and well-defined culturing protocol for LT-HSCs allowing for continuous expansion is lacking (Kumar and Geiger, 2017; Wilkinson et al., 2020). Production of LT-HSCs from iPSCs represents a promising alternative, however, differentiation protocols without ectopic transcription factor expression have not been established yet (Demirci et al., 2020). This may be due to the complex developmental programming of adult LT-HSCs that have experienced distinct niche signals during their migration from AGM via the fetal liver to the bone marrow, resulting in functional differences observed in fetal versus mature HSCs (Arora et al., 2014). Growing evidence, mainly provided by *in vitro* and *ex vivo* studies in combination with macrophage depletion via clodronate (Gao et al., 2021; Mariani et al., 2019), suggests that macrophages play an essential role in HSC development and maintenance during embryogenesis. However, it remains to be investigated whether specific macrophage populations in the fetal liver contribute to LT-HSC stem-cell ness or differentiation via factors they produce *in vivo*.

Given the observations implicating macrophages in HSC functionality, we characterized the heterogeneity and ontogeny of fetal liver macrophage populations, providing a comprehensive macrophage atlas of the fetal liver at E14.5. Further, using a conditional mouse model to deplete macrophages *in vivo*, we established fetal liver macrophages as important modulators of LT-HSC differentiation capacity into granulocytes.

## Results

### The fetal liver harbours heterogeneous macrophage populations

To investigate fetal liver macrophage heterogeneity, we performed single-cell RNA-sequencing on sorted CD11b^low/+^ cells at E14.5 (Figure S1A). To isolate macrophages and macrophage progenitors for further downstream analyses from this myeloid population, we clustered all cells and overlayed a pre-macrophage (pMac) signature (Mass et al., 2016). Out of eleven clusters, four clusters were chosen (Figure S1B, C, see Methods) and analysed further. To further select specifically macrophages, the same procedure was performed twice on the re-clustered cells using a signature enriched in fetal macrophages when compared with EMPs (Figure S1D), resulting in 18 clusters (Figure S1E). Cells that either expressed macrophage precursor genes (*Clec7a, Ccr2, Cx3cr1, Csf1r*), pan-macrophage genes (*Mrc1, Adrgre1, Siglec1, Msr1, Cd63*) and/or liver macrophage-specific genes (*Timd4, Clec4f, Vcam1*) were enriched in clusters 1, 2, 7, 8, 9, and 11, which were chosen for further downstream analysis (Figure 1A, B, S1F). The predominant expression of *Ly6c2, Ly6g, Cxcr2*, and *Cd33* in the remaining clusters (Figure S1F) indicated their monocyte/granulocyte or myelomonocytic precursor cell identity, respectively, and were therefore excluded from further analysis.

**Figure 1.**
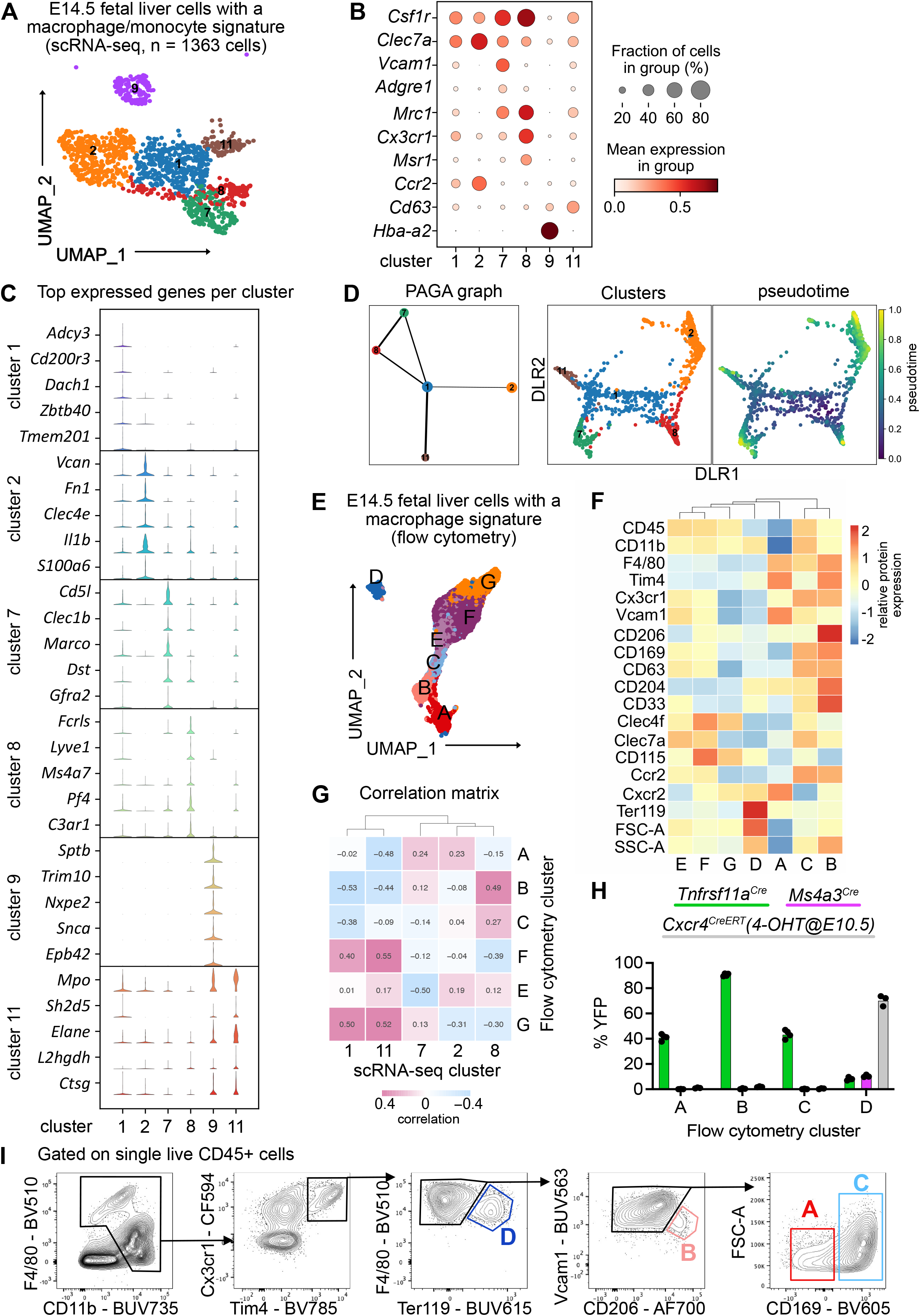
Characterization of fetal liver macrophage populations. **(A)** Single-cell RNA-sequencing (scRNA-seq) analysis of wildtype CD11b^low/+^ cells isolated from a fetal liver at developmental day (E)14.5. Clusters of possible macrophage subsets identified by a monocyte/macrophage signature (see Supplemental Figure 1) are visualized through UMAP. **(B)** Expression of selected macrophage- and macrophage progenitor-specific genes in clusters from (A). **(C)** Violin plots of highly expressed genes within the clusters from (A). **(D)** Developmental trajectory analysis using PAGA method (left) and pseudotime analysis (right) of the identified clusters in (A), excluding cluster 9. **(E)** Flow cytometry analysis of CD11b^low/+^ cells with a macrophage signature, isolated from a fetal liver at developmental day E14.5. Cell surface marker expression was used to generate unbiased clusters using UMAP. **(F)** Heatmap of relative protein expression and cell size parameters used in the flow-cytometry analysis in (E). **(G)** Correlation matrix between the flow cytometry and scRNA-seq data reveals highly correlating clusters between the two datasets. **(H)** Flow-cytometry analysis of E14.5 fetal liver macrophages using three different fate-mapping mouse models. CD11b^low/+^ cells with a macrophage signature were analyzed as shown in (E, F), resulting in similar cluster distribution (see Supplemental Figure 2C). YFP^+^ cells from the *Tnfrsf11a^Cre^; Rosa26^YFP^* model (green) indicate a yolk sac origin. *Ms4a3^Cre^; Rosa26^YFP^* (pink) and *Cxcr4^CreERT^; Rosa26^YFP^* with 4-hydroxytamoxifen (4-OHT) injection at E10.5 (grey) indicate a monocytic and hematopoietic stem cell origin, respectively. Circles represent individual mice. **(I)** Simplified gating strategy to identify macrophage clusters in E14.5 livers using flow cytometry.

Cluster 9 expressed almost exclusively erythroblast-specific genes (*Hba-a2, Sptb, Trim10, Nxpe2, Snca, Epb42*) (Figure 1B, C) and was identified as EI macrophages, which have been recently described as doublets of erythroblasts and macrophages (Popescu et al., 2019) or erythroblasts with cell remnants on their surface since macrophages frequently adhere to other cells (Millard et al., 2021). Clusters 7 and 8 showed the highest expression of bona fide macrophage genes such as *Csf1r, Mrc1*, and *Timd4* but were distinct in their expression of other macrophage markers (e.g., *Marco, Lyve1*) (Figure 1B, 1C, S1E). Cells in cluster 2 expressed the highest levels of *Ccr2, Clec7a, Clec4a, Il1b*, and *S100a6* (Figure 1B, C) besides some of the macrophage-specific genes, hinting towards an inflammatory state of this putative macrophage population. Clusters 1 and 11 showed a low expression of core macrophage genes compared to clusters 2, 7, and 8, suggesting a precursor stage (Figure 1B). However, cluster 11 was distinct from cluster 1 with a high expression of granule-related genes, such as *Mpo*, *Elane*, and *Ctsg* (Figure 1C), which may indicate a granulocytic rather than a macrophage precursor state. To test these hypotheses and predict developmental trajectories, we performed a Partition-based graph abstraction (PAGA) analysis (Wolf et al., 2019) after excluding cluster 9 due to their doublet identity. Here, cluster 1 expression represents a progenitor state, thus, is the centre of the network, which was confirmed by pseudotime analysis (Figure 1D). The other clusters fall into distinct nodes, with clusters 7 and 8 showing substantial similarity as indicated by the edge thickness (Figure 1D). In summary, our scRNA-seq analysis indicates the presence of at least three macrophage states (clusters 2, 7, and 8), which have distinct phenotypes.

To validate macrophage heterogeneity on the protein level, we performed a high-dimensional flow cytometry analysis on CD11b^low/+^ cells (Figure S2A). Similar to the scRNA-seq enrichment, we visualised all myeloid cells using UMAP, clustered them, and overlayed a macrophage signature (F4/80, Tim4, Cx3cr1, Vcam1, CD169, CD206, Figure S2B). This resulted in seven clusters, which we analysed further (Figure 1E). Hierarchical clustering of all clusters expressing macrophage and macrophage precursor proteins confirmed the presence of three F4/80^high^ macrophage populations (clusters A-C, Figure 1F). As already observed on the transcriptional level, cluster D likely represents Ter119^+^ erythroblasts with macrophage cell remnants, as indicated by the increased cell size and granularity determined via FSC and SSC, respectively (Figure 1F). To correlate precursor and macrophage clusters identified by transcriptional and protein analyses, a correlation matrix between the scRNA-seq and flow cytometry data sets was calculated based on gene expression corresponding to the presence of the respective antigen used in our flow cytometry panel (Figure 1G). Here, scRNA-seq precursor clusters 1 and 11 corresponded highly to clusters F and G, which expressed high levels of CD45, CD115 and Clec4f but were low in F4/80 and Tim4 expression. In contrast, scRNA-seq clusters 2 and 7 represented cluster A while cluster 8 correlated mostly with clusters B and C, supporting the notion of three distinct macrophage populations in the fetal liver at E14.5, in addition to EI macrophages.

### Fetal liver macrophages originate from yolk-sac progenitors

Next, we addressed the ontogeny of the macrophage clusters using *Rosa26^LSL-YFP^* mice crossed to *Tnfrsf11a^Cre^* for detection of pMac-derived cells (Mass et al., 2016), to *Ms4a3^Cre^* for detection of monocyte-derived cells (Liu et al., 2019) and to the inducible *Cxcr4^CreERT^* with 4-hydroxytamoxifen injection at E10.5 labelling all cells of the definitive hematopoiesis wave (Werner et al., 2020). We confirmed the presence of all three macrophage clusters and the Ter119^+^ EI macrophage cluster in all mouse models (Figure S2C). HSC-derived definitive erythroblasts in cluster D were efficiently fate-mapped using the *Cxcr4^CreERT^* model, validating that these were cell doublets or erythroblasts with cell remnants on their surface since they also showed low labelling of the other fate-mapping models that label macrophages/monocytes (Figure 1H). The remaining clusters were YFP^+^ only in the *Tnfrsf11a^Cre^* model, demonstrating that all fetal liver macrophages at E14.5 derive from pMacs. Using the distinct expression of Vcam1, CD206 and CD169 in clusters A-C and their difference in cell size allowed us to develop a simple gating strategy to distinguish these macrophage populations (Figure 1I). In summary, using a hypothesis-driven analysis of CD11b^low/+^ cells, we define, in addition to the already well-known EI macrophage population, distinct macrophage populations in the fetal liver that are yolk sac-derived.

### Fetal liver macrophage subpopulations display distinct transcriptional programs

Next, we set out to investigate whether the macrophage heterogeneity we defined at E14.5 could indicate additional macrophage states besides serving as EI macrophages, therefore, resulting in other functions than phagocytosing erythroblast nuclei. To this end, we first performed a GO term analysis on the top expressed 100 genes, comparing each cluster from the scRNA-seq analysis to all other clusters. Here, cluster 2-specific genes fell into the terms ‘cell chemotaxis’, ‘positive regulation of inflammatory response’, and ‘cytokine-mediated signaling pathway’ (Figure 2A), indicating the activated inflammatory state already observed in the top five expressed genes (Figure 1C). In contrast, cluster 7-related genes were significant for the GO terms ‘positive regulation of lipid localization’, ‘ameboidal-type cell migration’, and ‘apoptotic cell clearance’. Also cluster 8 expressed genes belonging to the GO term “positive regulation of lipid localization’ that partially overlapped with cluster 7 (e.g., *Apoe, Lpl*) (Table S1). Additional cluster 8-specific terms were ‘macrophage activation’ and ‘cell junction disassembly’ (Figure 2A). This analysis indicated that, albeit somewhat similar, the distinct macrophage populations might exert distinct functions in the fetal liver. This was confirmed by the cluster-specific expression of selected ligands indicating that the distinct cellular states may result in distinct paracrine signalling activity (Figure 2B). Intersecting the CellTalk database information (Shao et al., 2021) with the complete set of genes expressed by the three macrophage states revealed 208 potential macrophage-derived secreted ligands (Table S2).

**Figure 2.**
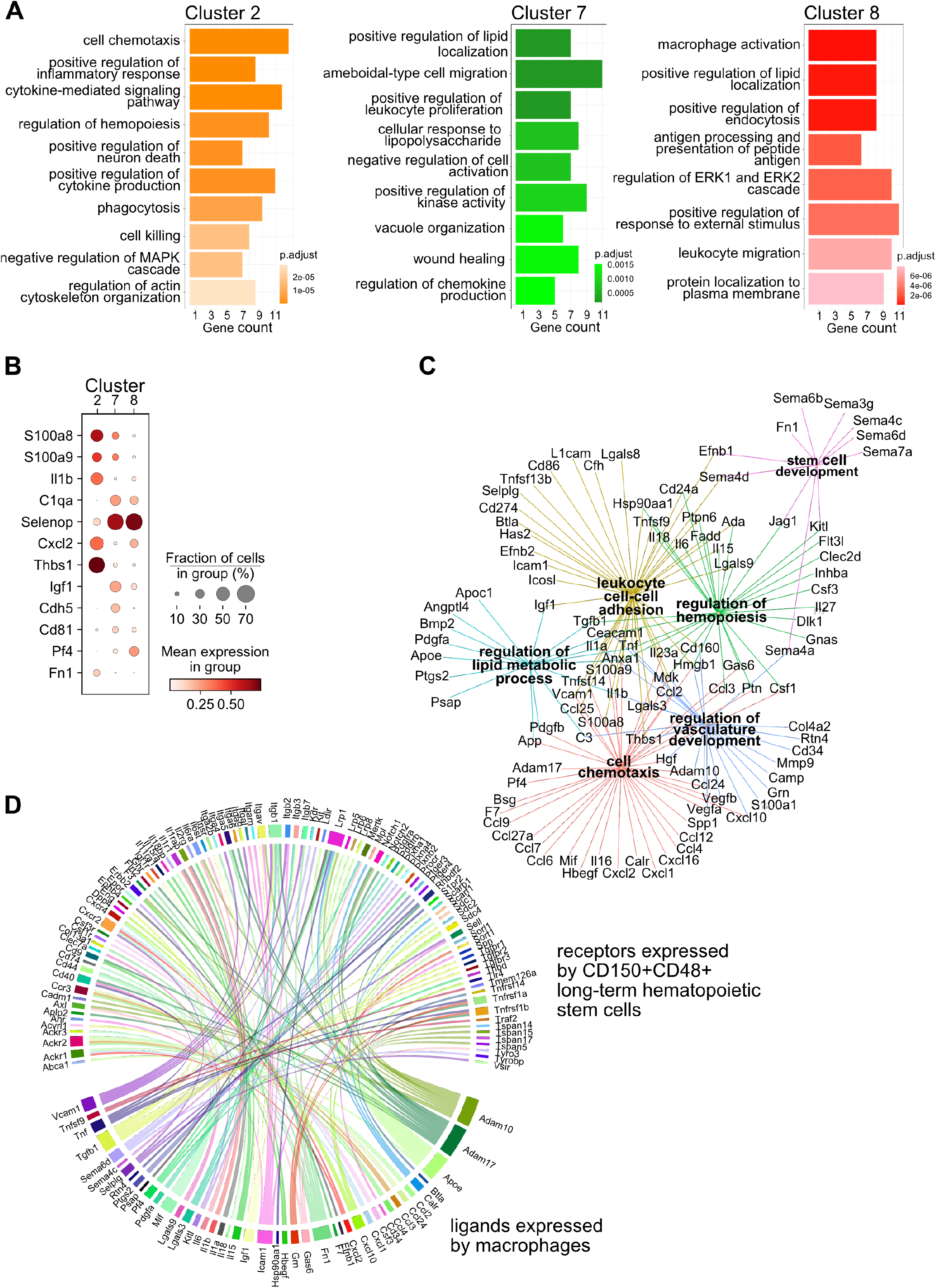
Transcriptional program and paracrine signalling of fetal liver macrophages. **(A)** Gene set enrichment analysis (GSEA) of final macrophage clusters 2, 7 and 8 was performed on the differentially expressed genes of each cluster. **(B)** Expression of selected ligands in the final macrophage clusters. **(C)** Interaction and functional network of expressed ligands on the identified macrophage clusters 2, 7 and 8. Each hub with a colour indicates the function of the ligands, while the edges show the interaction between them. **(D)** Potential ligand-receptor interactions between macrophages and LT-HSCs. The gene names on the bottom of the plot are expressed ligands in macrophage clusters 2, 7 and 8. Gene names on the top are expressed receptors on LT-HSCs at E14.5. Each ligand can target several receptors which are indicated with the same colour.

We next used the 208 ligand candidates and performed a Gene Set Enrichment Analysis, and visualized the gene/GO-term relationships in a network (Figure 2C). This analysis pointed to additional functionality of the three macrophage states beyond erythropoiesis, which included regulation of hemopoiesis and stem cell development together with chemotaxis and vasculature development, mechanisms that could shape the stem cell niche.

Macrophages contribute to the stem cell niche, particularly in the bone marrow, under inflammatory conditions (Seyfried et al., 2020). Yet, evidence for macrophage-derived molecules involved in the direct cell crosstalk controlling stem cell maintenance and differentiation in the mouse fetal liver is missing. Therefore, we sought to explore potential signalling events between macrophages and HSCs to determine whether macrophage-derived factors might modify HSC function during steady-state. Thus, we sequenced LT-HSC at E14.5 and, leveraging our scRNA-seq macrophage dataset (see Methods), uncovered potential ligand-receptor interactions between macrophage-derived ligands and LT-HSCs in the fetal liver (Figure 2D). Many of the ligands are well-known players in the stem cell niche, e.g., Kitl, Igf1, Tnf, Tgfb1, and Fn1, which have been reported to directly or indirectly promote the expansion of hematopoietic stem and progenitor cells (Azzoni et al., 2018; Hadland et al., 2022; Sakaki-Yumoto et al., 2013). In summary, our transcriptomic ligand-receptor interaction analyses suggest that macrophages express HSC niche factors and, thereby, may actively contribute to LT-HSC functionality in the fetal liver.

### Macrophages interact directly with LT-HSCs

We next asked whether the potential paracrine signalling between macrophages and LT-HSCs in the fetal liver occurs via direct interaction. To test this, we first performed 3D whole-mount immunofluorescence analyses in E14.5 livers. We found EI macrophages that were entirely surrounded by Ter119^+^ erythroblasts, as expected, but frequently observed cell-cell interactions of macrophages with c-Kit^+^ progenitors and CD150^+^ LT-HSCs (Figure 3A). To determine whether LT-HSCs interact with distinct macrophage populations, we assessed cellular interactions via co-detection by indexing (CODEX)-enabled high-dimensional imaging (Black et al., 2021; Frede et al., 2022; Goltsev et al., 2018), validating the presence of clusters A-D identified by flow cytometry. CD45, Iba1, F4/80, Cx3cr1 and Tim4 were used as pan-macrophage markers (Figure 3B), allowing the distinction of cluster A: CD106 (Vcam1)^+^ macrophages, cluster B: CD206^+^ macrophages, cluster C: CD169^+^ macrophages and the most abundant cluster D: Ter119^+^ EI macrophages (Figure 3B, Figure S3A). First, we assessed the distribution of the four macrophage clusters in the different liver lobes using a Voronoi diagram, showing that all macrophages are found across the whole tissue (Figure 3C). CD150^+^ HSC were also dispersed across the fetal liver but showed a preferential localization near the liver capsule (Figure 3C). Similar results were detected via spatially segmented cellular neighbourhoods of the single objects using a raster scan with a radius of 50 μm and a self-organizing map (SOM) algorithm where CD150^+^ HSCs, represented by neighbourhood 4, were mainly found near the liver capsules of the lower left and the two upper liver lobes (Figure S3B). Next, we used a data-driven approach to detect spatial interactions between macrophage clusters and HSCs within a range of 5 to 50 μm. Interestingly, LT-HSCs showed the highest correlation with macrophages from cluster C representing CD169^+^ macrophages (Figure 3D) and the lowest correlation with cluster D (Ter119^+^ macrophages). Manual inspection of cells surrounding CD150^+^ HSC in a 50 μm radius also revealed that LT-HSCs were preferentially surrounded by CD169^+^ macrophages (cluster C, Figure 3E), with fewer identified cells belonging to the other macrophage populations (Figure 3F).

**Figure 3.**
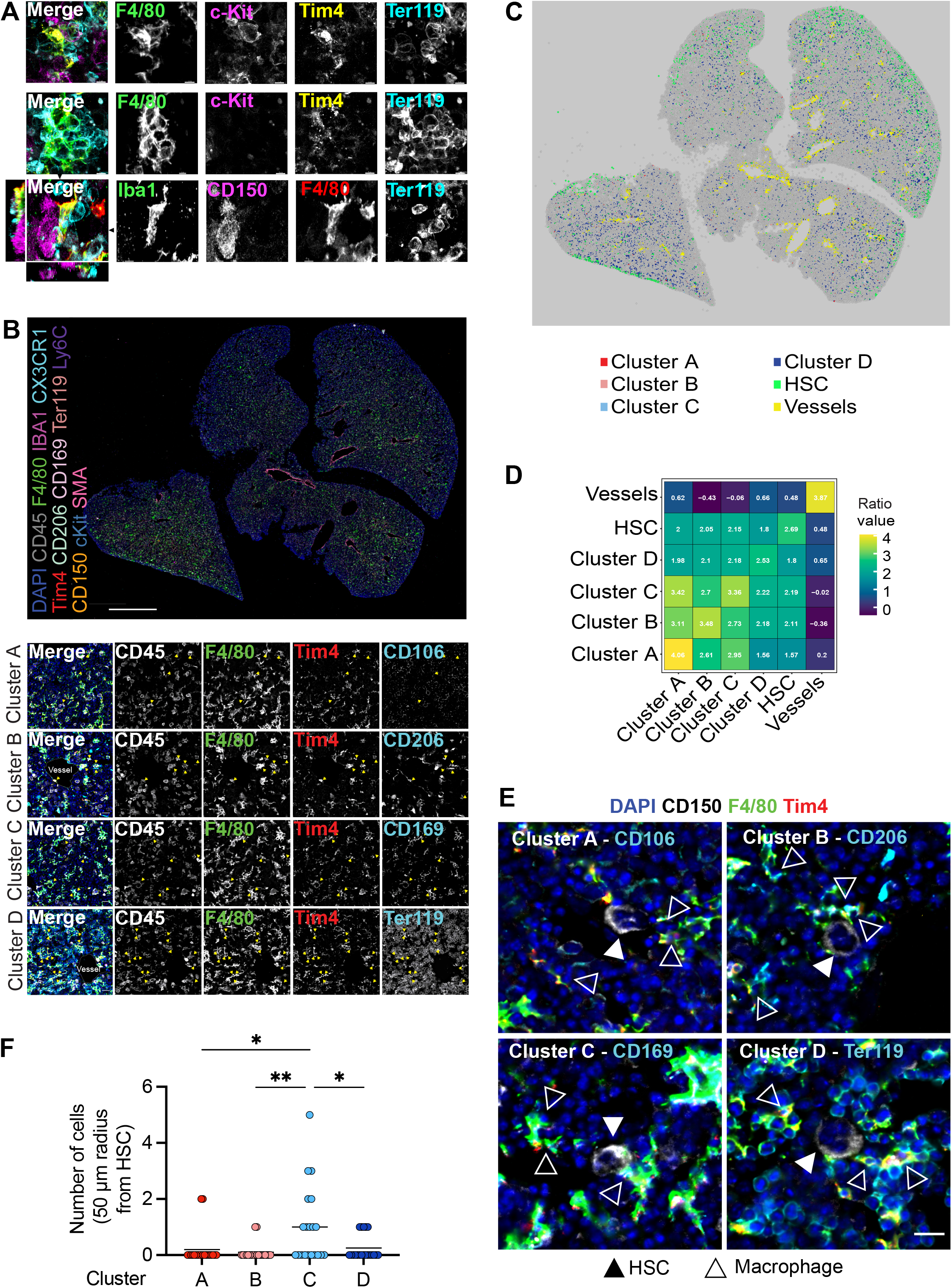
Macrophage heterogeneity and their interaction with HSCs. **(A)** Immunostaining of E14.5 fetal liver sections with antibodies against F4/80, c-Kit, Tim4, Ter119, and CD150. The first row visualizes the interaction between EI macrophages surrounded by Ter119^+^ erythroblasts. The second and last rows highlight the interaction between macrophages and progenitor cells, including CD150^+^ LT-HSCs. The black arrows on the last row indicate the yz and xz dimensions which are shown on the left and bottom sides. **(B)** A 5 μm frozen section of a fetal liver from an E14.5 wildtype embryo was stained with a 20-plex CODEX antibody panel. Representative image of the entire field of view is shown in the upper panel. Enlargements showing representative images of the cells from clusters A-D are shown in the lower panel. Yellow arrows in the enlargements indicate macrophages from the corresponding cluster. Scale bars represent 500 μm in the overlay and 15 μm in the enlargements. **(C)** Voronoi diagram from (B) after manual cell classification using the HSCs, blood vessels, and cells from the four macrophage clusters as seeds. **(D)** Spatial analysis of interactions between cells from macrophage clusters, HSCs, and blood vessels in the fetal liver within a range of 5 to 50 μm. Values represent the calculated Log Odds Ratio. **(E)** Representative images of the interaction of HSCs with the macrophage clusters. Filled arrowheads indicate the HSC, and empty arrowheads indicate the macrophages from the corresponding cluster. Scale bar represents 15 μm. **(F)** Absolute number of cells from the four macrophage clusters within a radius of 50 μm from one randomly selected HSC. Each dot represents one cell. Black lines in the plot represent the mean. One-way ANOVA; * *p* < 0.05, ** *p* < 0.01.

As erythroblasts are the most abundant cell type, we asked whether clusters A-C serve, at least partially, as EI macrophages. To this end, 50 macrophages of each cluster were randomly chosen, and the direct interaction with Ter119^+^ cells was evaluated manually. While cells belonging to cluster D (Ter119^+^ EI macrophages) showed 100 % interaction, as expected, cluster A (CD106^+^ macrophages, 73 % interaction), cluster B (CD206^+^ macrophages, 80 % interaction), and cluster C (CD169^+^ macrophages, 80.4 % interaction) did not always interact with Ter119^+^ erythroblasts (Figure S3C, D). The CD206^+^ cluster A was the subpopulation with the least interaction and longest distance to the nearest erythroblast (Figure S3E), which was often accompanied by an elongated cell shape near vessels, indicating the presence of CD206^+^ perivascular macrophages (Figure 3B). Of note, the tissue analysed via CODEX represents only cellular neighbourhoods in X and Y due to the thin sectioning technique (5 μm) and, thus, does not take neighbouring cells in the Z plane into account. This leads to an underestimation of macrophage-HSC interactions, as indicated by F4/80^+^ filopodia extending towards CD150^+^ cells in almost all cases (Figure 3E). In summary, our data indicate that macrophages inhabit distinct niches within the fetal liver, with the majority of macrophages supporting mainly erythroblast maturation, while other populations may support HSC function.

### Lack of macrophages leads to decreased erythrocyte maturation

Given that LT-HSCs have direct contact with or are in close proximity to macrophages providing niche signals, we hypothesised that depletion of fetal liver macrophages would alter the LT-HSC phenotype and function. Therefore, we took advantage of the *Tnfrsf11a^Cre/+^; Spi1^f/f^* mouse model, which should lead to a depletion of fetal macrophages since *Tnfrsf11a* is expressed by pMacs (Mass et al., 2016) and *Spi1* (also known as *Pu.1*) is required for macrophage differentiation (McKercher et al., 1996; Scott et al., 1994). Indeed, flow cytometry and immunostaining revealed an 80-90 % reduction of F4/80^+^/Iba1^+^ cells in fetal livers of *Tnfrsf11a^Cre/+^; Spi1^f/f^* embryos compared to littermate controls demonstrating efficient depletion of macrophages (Figure 4A, B). First, we analysed the absolute cell numbers and the number of CD45^+^ cells per fetal liver (Figure 4C), to ensure that our depletion strategy did not have any major off-targets leading to a developmental delay. The lack of an overall change of tissue architecture was confirmed by a haematoxylin-eosin staining (Figure 4D). Erythroblast maturation is characterized by changes in CD71 and Ter119 expression, allowing us to determine their developmental sequence within six subsets (from S0 to S5) (Fraser et al., 2007; Pop et al., 2010). The final maturation step of enucleation to produce a functional erythrocyte depends on EI macrophages (Palis, 2014). Indeed, Giemsa staining of blood smear samples from *Tnfrsf11a^Cre/+^; Spi1^f/f^* knockout embryos showed a reduction of enucleated erythrocytes compared to controls (Figure 4E). However, the maturation of erythroblasts was not altered (Figure 4F). These results suggest that our newly developed mouse model efficiently targets fetal liver macrophages at E14.5, leading to a delayed erythrocyte enucleation, consistent with the function of EI macrophages.

**Figure 4.**
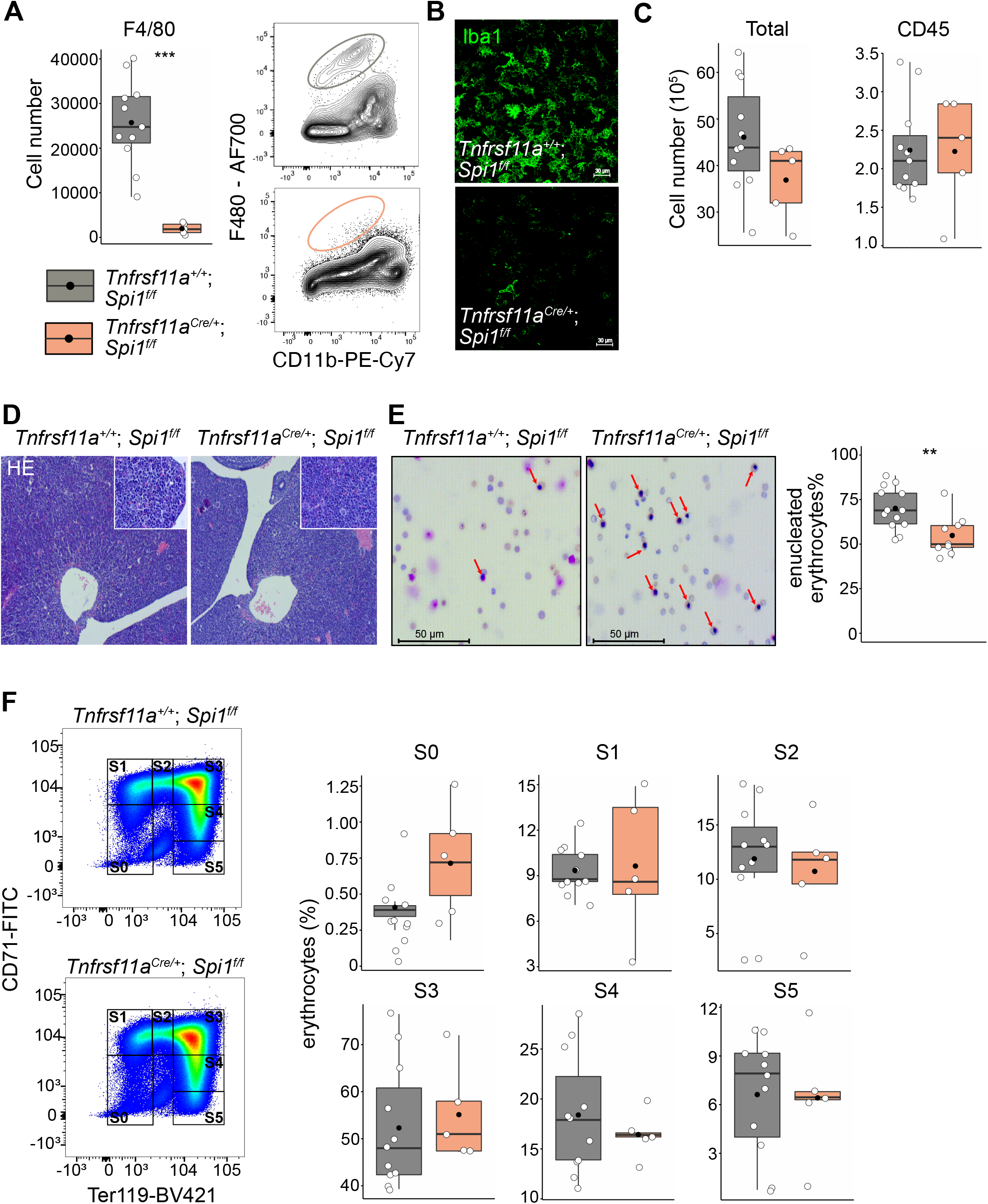
The effect of macrophage depletion on erythropoiesis. **(A)** Quantification of F4/80^+^ macrophage cells in the E14.5 fetal liver of control and *Tnfrsf11a^Cre/+^; Spi1^f/f^* knockout embryos using flow cytometry. n = 11 for *Tnfrsf11a^+/+^; Spi1^f/f^*, n = 5 *Tnfrsf11a^Cre/+^; Spi1^f/f^*. **(B)** Immunofluorescent staining of macrophages using Iba1 antibody in E14.5 livers. Representative for n = 3. Scale bar represents 30 μm. **(C)** Quantification of single live and CD45^+^ cells in the E14.5 fetal liver of control and *Tnfrsf11a^Cre/+^*; *Spi1^f/f^* knockout embryos using flow cytometry. n = 11 for *Tnfrsf11a^+/+^; Spi1^f/f^*, n =5 *Tnfrsf11a^Cre/+^; Spi1^f/f^*. (**D**) Haematoxylin and eosin stain (HE) of *Tnfrsf11a^+/+^*; *Spi1^f/f^* and *Tnfrsf11a^Cre/+^*; *Spi1^f/f^* fetal livers at E14.5. Representative for n = 5. Overviews were taken with a 5x objective, insets with a 20x objective. **(E)** On the left: representative pictures of blood smear using May-Grünwald-Giemsa staining. Arrows indicate nucleated erythroblasts. On the right; the percentage of enucleated erythrocytes in the blood of *Tnfrsf11a^+/+^; Spi1^f/f^* and *Tnfrsf11a^Cre/+^; Spi1^f/f^* embryos at E14.5. n =13 for *Tnfrsf11a^+/+^; Spi1^f/f^* and n = 9 for *Tnfrsf11a^Cre/+^; Spi1^f/f^*. **(F)** On the left: representative gating strategy to capture differentiation stages of erythrocytes. On the right: comparison of the erythrocyte’s percentages in each of the differentiation stages between *Tnfrsf11a^+/+^; Spi1^f/f^* and *Tnfrsf11a^Cre/+^; Spi1^f/f^* embryos at E14.5. n =11 for *Tnfrsf11a^+/+^; Spi1^f/f^* and n = 5 *forTnfrsf11a^Cre/+^; Spi1^f/f^*. All statistical tests comparing control and *Tnfrsf11a^Cre/+^; Spi1^f/f^* embryos: ****P<0.0001, ***P≪0.001, **P<0.01, and *P<0.05, Wilcoxon test.

### Depletion of macrophages leads to transcriptional changes in HSCs

To determine whether the lack of macrophages would impact LT-HSC functionality, we performed bulk RNA-sequencing on sorted LT-HSCs from *Tnfrsf11a^Cre/+^; Spi1^f/f^* knockout embryos and littermate controls at E14.5 (Figure S4A). Analysis of differentially expressed genes (DEG) resulted in 598 upregulated and 555 downregulated genes (Figure 5A, S4B, Table S3). Some of the upregulated genes were well-known transcriptional regulators of hematopoietic specification and stem cell capacity, such as *Gata2* and *Gata3*. (Figure 5A, S4C). Examining Gene Ontology (GO) pathways of these DEG revealed signalling mechanisms enriched for metabolic processes, organelle localization and RNA-related processes to be downregulated (Figure 5B, Table S4). In contrast, genes belonging to the GO terms chromatin organization, myeloid cell differentiation, regulation of hemopoiesis and mononuclear cell proliferation were upregulated (Figure 5B, S4C). These data indicate that the transcriptional program of LT-HSC is regulated by macrophages and, thereby, may impact their proliferative and/or differentiation potential.

**Figure 5.**
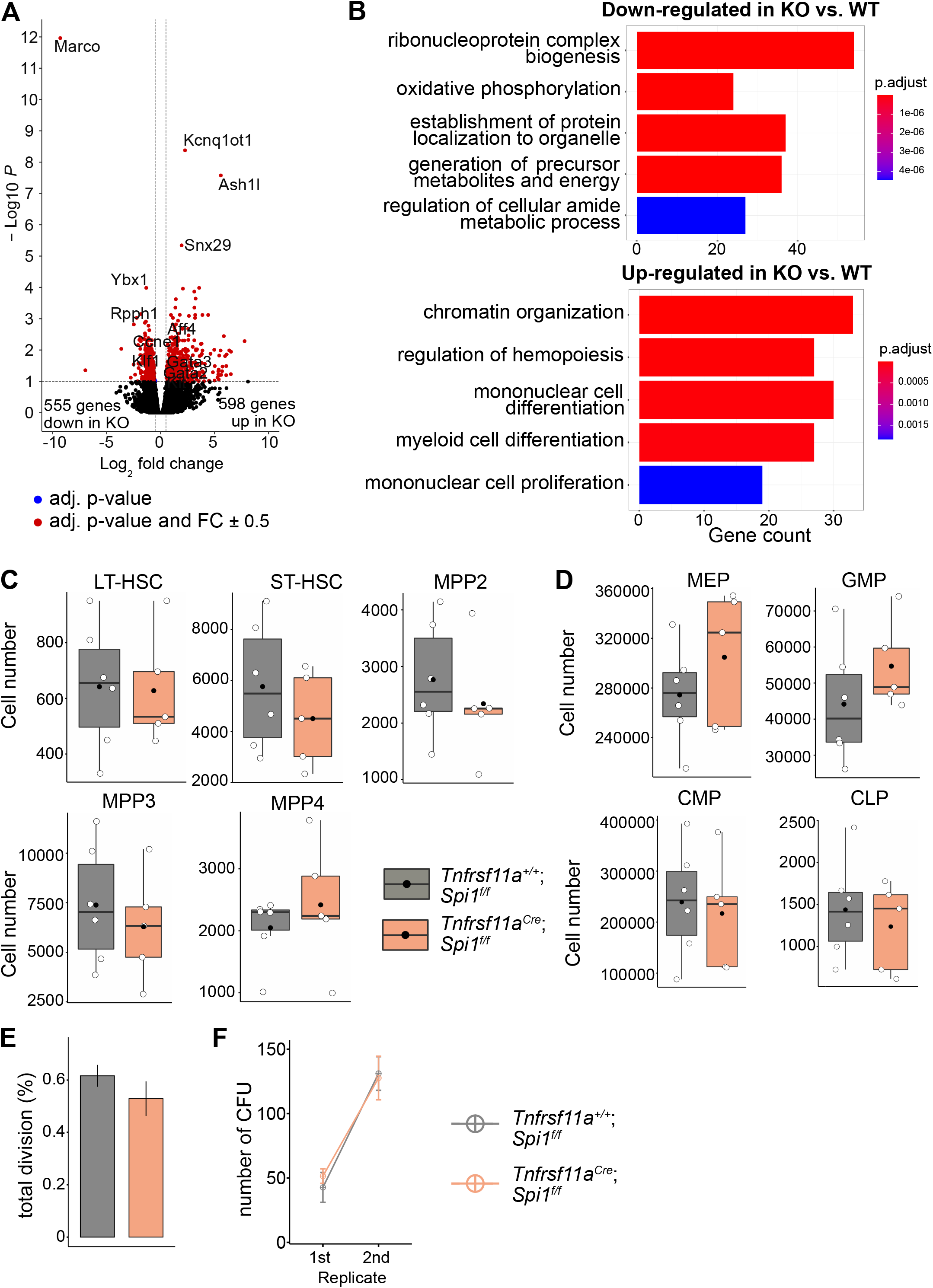
The effect of macrophage depletion on haematopoiesis. **(A)** Volcano plot of differentially expressed genes between CD150^+^ LT-HSCs sorted from *Tnfrsf11a^+/+^; Spi1^f/f^* (n = 4) and *Tnfrsf11a^Cre/+^; Spi1^f/f^* (n = 6) embryos. Blue dots are significant (adjusted p-value <0.1), red dots are significant with a fold-change of ± 0.5. **(B)** Gene set enrichment analysis of significant up and down-regulated genes from (A) between the control and knockout embryos. **(C)** Quantification of the stem- and progenitor cells from *Tnfrsf11a^+/+^; Spi1^f/f^* and *Tnfrsf11a^Cre/+^; Spi1^f/f^* fetal livers at E14.5. LT-HSC: long-term hematopoietic stem cells; ST-HSC: short-term hematopoietic stem cells; MPP: multipotent progenitors. n = 6 for *Tnfrsf11a^+/+^; Spi1^f/f^* and n = 5 *forTnfrsf11a^Cre/+^; Spi1^f/f^*. **(D)** Quantification of progenitors from *Tnfrsf11a^+/+^; Spi1^f/f^* and *Tnfrsf11a^Cre/+^; Spi1^f/f^* fetal livers at E14.5. n = 6 for *Tnfrsf11a^+/+^; Spi1^f/f^* and n = 5 for *Tnfrsf11a^Cre/+^; Spi1^f/f^*. CLP: common lymphoid progenitor; CMP: common myeloid progenitor; GMP: granulocyte-macrophage progenitor; MEP: megakaryocyte-erythrocyte progenitor. **(E)** Cell proliferation assay. LT-HSCs were harvested from the fetal liver at E14.5 using FACS for performing a single-cell colony assay. n = 77 LT-HSCs from n = 5 fetal livers for *Tnfrsf11a^+/+^; Spi1^f/f^* and n = 96 LT-HSCs from n = 5 fetal livers *forTnfrsf11a^Cre/+^; Spi1^f/f^*. **(F)** Serial transfer colony-forming assay showing the number of observed colonies after seeding E14.5 fetal liver cells into media (1^st^ replicate) and re-seeding the cultured colonies (2^nd^ replicate). n = 7 for *Tnfrsf11a^+/+^; Spi1^f/f^* and n = 8 for*Tnfrsf11a^Cre/+^*; *Spi1^f/f^*.

### Depletion of macrophages does not change stem and progenitor cell numbers

To assess LT-HSC maintenance, retention and proliferation in the fetal liver, we first performed flow cytometry experiments to quantify stem and progenitor cell numbers (gating strategy Figure S5A). Quantification of LT-HSCs, short-term (ST)-HSCs, multipotent progenitors (MPP) 2, MPP3, and MPP4 did not reveal any significant differences between *Tnfrsf11a^Cre/+^*; *Spi1^f/f^* knockout embryos and littermate controls at E14.5 (Figure 5C). Further downstream progenitors, such as the common lymphoid progenitor (CLP), common myeloid progenitor (CMP), megakaryocyte-erythroid progenitor (MEP) and granulocyte-macrophage progenitor (GMP), were not significantly altered in cell numbers, albeit there was a tendency for increased GMP numbers in *Tnfrsf11a^Cre/+^; Spi1^f/f^* knockout livers (Figure 5D). To address the proliferation capacity of HSCs, we sorted single LT-HSCs from *Tnfrsf11a^Cre/+^*; *Spi1^f/f^* and littermate controls and monitored their proliferation 48 hours later (Figure 5E). Further, we performed serial colony-forming unit (CFU) assays to study the long-term self-renewal ability (Figure 5F). In both assays, *Tnfrsf11a^Cre/+^; Spi1^f/f^* stem cells showed no defects or increase in proliferation compared to littermate controls (Figure 5C, D). These results suggest that a reduction of macrophages in the HSC niche does not modify stem and progenitor cell numbers or lead to a dysregulated proliferation capacity of LT-HSCs.

### Macrophages control HSC differentiation potential

Next, we addressed the differentiation behaviour of HSCs *in vivo* and *in vitro*. Using flow cytometry, we focused on myeloid cells since genes important for myeloid cell differentiation were upregulated (Figure 5B, S4C). An unbiased clustering of cells using UMAP indicated a reduction of F4/80^+^ macrophage clusters A-C, but not of cluster D (Figure 6A), which again underlines the fact that these F4/80^+^Ter119^+^ events represent cell doublets or erythroblasts with attached macrophage cell remnants. In addition to a Ly6G^+^ neutrophil cluster, we detected two Ly6C^+^ monocyte clusters that were distinguished by their Cx3cr1 expression. Using a gating strategy to detect these myeloid cell types (Figure S5B), we observed a significant increase of Ly6G^+^ cells in *Tnfrsf11a^Cre/+^; Spi1^f/f^* compared to littermate controls while the number of monocytes was not altered (Figure 6B). To test whether the change in differentiation potential of *Tnfrsf11a^Cre/+^*; *Spi1^f/f^* HSCs is cell-autonomous, we performed a CFU-C assay. While the total number of colonies was similar in both genotypes, the number of CFU-granulocyte/macrophage (GM) was increased (Figure 6C). Together with the results from the RNA-seq analyses, our data indicate that the lack of fetal liver macrophages causes a reprogramming of LT-HSCs, leading to their preferential differentiation towards the granulocytic lineage.

**Figure 6.**
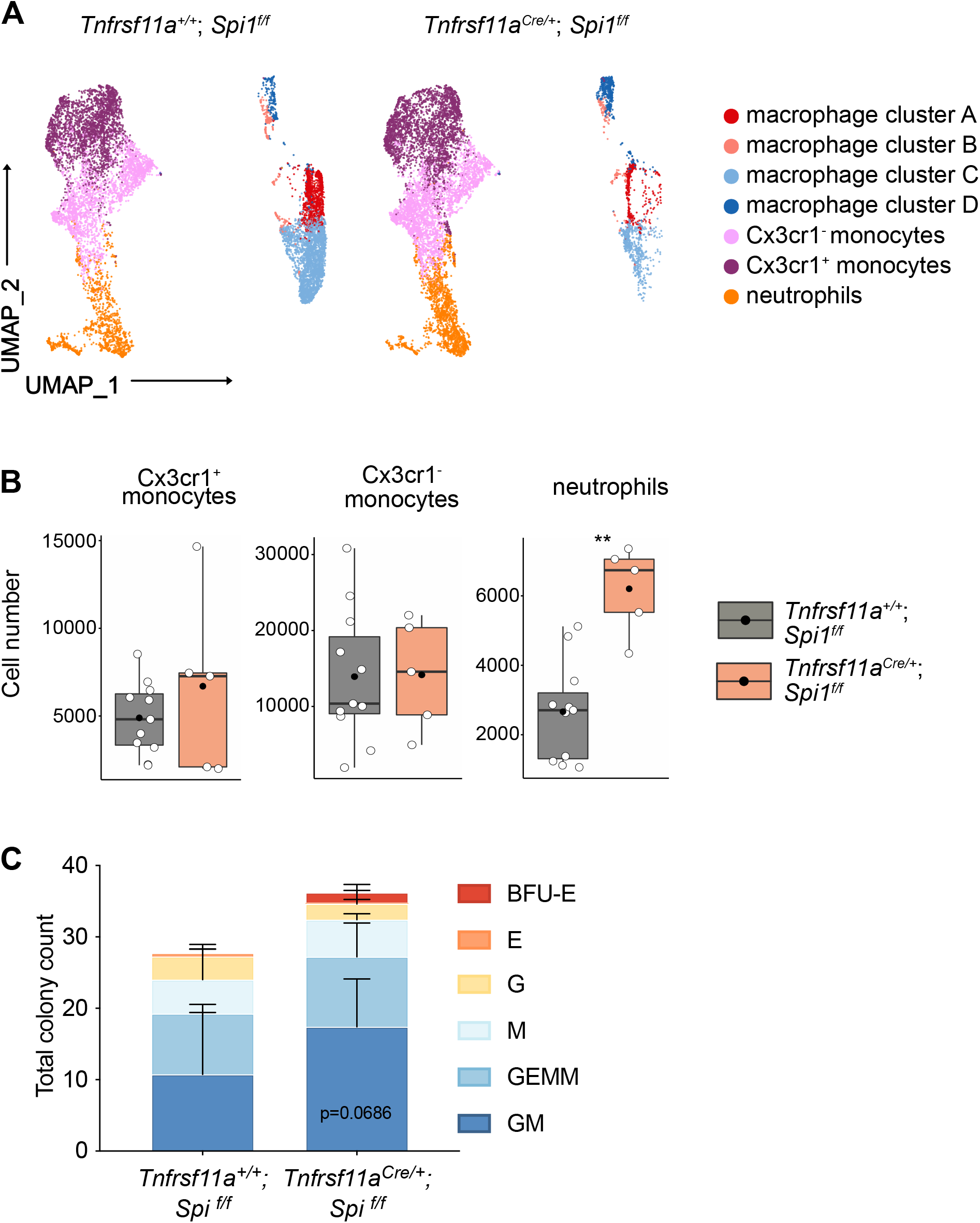
Macrophage depletion shifts haematopoiesis towards the granulocytic lineage. **(A)** Flow cytometry analysis of CD11b^low/+^ cells isolated from *Tnfrsf11a^+/+^; Spi1^f/f^* and *Tnfrsf11a^Cre/+^; Spi1^f/f^* fetal livers at E14.5. Cell surface marker expression was used to generate unbiased clusters using UMAP, which were subsequently used for a gating strategy to quantify respective populations. **(B)** Quantification of monocytes (Cx3cr1^+^ and Cx3cr1^-^) and neutrophils progenitors from *Tnfrsf11a^+/+^; Spi1^f/f^* and *Tnfrsf11a^Cre/+^; Spi1^f/f^* fetal livers at E14.5. n = 11 for *Tnfrsf11a^+/+^; Spi1^f/f^* and n = 5 *forTnfrsf11a^Cre/+^; Spi1^f/f^*. **(C)** Colony-forming unit assay from *Tnfrsf11a^+/+^; Spi1^f/f^* and *Tnfrsf11a^Cre/+^; Spi1^f/f^* fetal livers at E14.5. n = 4 for *Tnfrsf11a^+/+^; Spi1^f/f^* and n = 5 *forTnfrsf11a^Cre/+^; Spi1^f/f^*. 2way ANOVA test. BFU: Burst-forming erythroid, E: erythroid; G granulocyte; GEMM; granulocyte, erythroid, macrophage, megakaryocyte; GM granulocyte, macrophage; M: macrophage.

## Discussion

We have shown that liver macrophages at E14.5 are heterogenous and that they play an active role in the niche of CD150^+^ LT-HSCs. While most macrophages serve purely as EI macrophages, being only surrounded by Ter119^+^ erythroblasts and promoting erythroblast maturation, a subset of macrophages directly interacts with other cells, such as c-Kit^+^ and CD150^+^ stem and progenitor cells. Being part of the HSCs niche, macrophages seem to specifically control the production of neutrophils, likely via paracrine factors that imprint the tissue environment on LT-HSCs, thereby enabling a tight balance of hematopoietic cell numbers. Using various fate-mapping models, we could exclude any contribution of definitive hematopoietic stem cells to the fetal liver macrophage pool. Finally, we provide a simple gating strategy with commonly available antibodies that allow the identification of macrophage subpopulations and the discrimination from cell doublets.

CODEX analyses indicate that CD150^+^ LT-HSCs are preferentially found in close proximity to CD169^+^ macrophages. CD169, also known as sialoadhesin, is a cell adhesion protein that has been described as an EI macrophage marker in the bone marrow and the fetal liver (Chow et al., 2013, 2011; Li et al., 2019; Seu et al., 2017), albeit with varying expression patterns (Seu et al., 2017). In line with this, our flow cytometry and *in situ* immunofluorescent CODEX analysis of the fetal liver indicates that the majority of macrophages express CD169, with few macrophages that are CD169-negative and small in size but that express high levels of F4/80, Tim4 and Vcam1, and can thus be considered *bona fide* macrophages (cluster A). CD169 has been further defined as essential for erythropoiesis by promoting erythroblast maturation in the bone marrow (Chow et al., 2013). Intriguingly, CD169 is not required to bind erythroblasts, as shown by studies using specific inhibitors (Morris et al., 1991, 1988), but instead accumulates in contact zones between macrophages and immature granulocytes (Crocker et al., 1990). However, these studies, as many others analysing EI macrophages, were performed after flushing the bone marrow. Thus, an ultrastructural characterization of the fetal liver *in situ* may be helpful in addressing whether CD169 forms clusters in the plasma membrane, which may be a direct interaction zone between CD169^+^ macrophages and LT-HSCs. However, since CD150^+^ LT-HSCs also interacted with CD169^-^ macrophages, the tethering mechanism may rely on another surface receptor altogether.

Recent studies have addressed the role of fetal macrophages in the development of hematopoietic stem and progenitor cells. Work in zebrafish shows a homing and retention mechanism controlled by macrophages (Li et al., 2018; Theodore et al., 2017). Li et al. describe a Vcam1^+^ macrophage-like cell population that interacts with hematopoietic stem and progenitor cells (HSPCs) and serves as a permissive signal for HSPC entry into the embryonic caudal haematopoietic tissue (CHT) niche (Li et al., 2018). Yet, our work does not support an essential role of macrophages for LT-HSC homing or retention to the fetal liver since numbers of LT-HSC are not affected in *Tnfrsf11a^Cre/+^; Spi1^f/f^* embryos. Work in mouse embryos indicates that CD206^+^ macrophages in the AGM contribute to intra-aortic HSC generation and maturation (Mariani et al., 2019). Further, macrophages have been suggested to promote HSC/MPP proliferation in the fetal liver (Gao et al., 2021). However, due to the lack of genetic mouse models targeting only yolk sac-derived macrophages and not the definitive wave of hematopoiesis, these studies relied on macrophage depletion via clodronate liposomes and the Csf1r inhibitor BLZ945 (Gao et al., 2021; Mariani et al., 2019). Thus, the long-term impact of these substances on the proliferation capacity of HSCs or other niche cells that promote HSC development cannot be excluded. Indeed, our model, in which we target pMacs very efficiently using the *Tnfrsf11a^Cre^* mouse model (Mass et al., 2016), leading to an almost complete depletion of macrophages while HSCs remain wildtype, we do not observe any defects in the proliferation and expansion of HSCs arguing for an unspecific off-target effect of clodronate and BLZ945.

During steady-state adulthood, macrophages have been shown to control neutrophil numbers through the clearance of apoptotic neutrophils and via the G-CSF/IL-17/IL-23 cytokine axis, which promotes granulopoiesis (Gordy et al., 2011; Hong et al., 2012; Stark et al., 2005). The reduction of macrophage populations in the bone marrow and spleen observed in *LysM^Cre^; c-Flip^f/f^* mice led to neutrophilia during steady state, which was attributed to the defect in efferocytosis of apoptotic neutrophils (Gordy et al., 2011). Furthermore, the *LysMC^re^; c-Flip^f/f^* model is also defined by an increase of inflammatory monocytes in the blood and spleen, indicating an alteration of myeloid progenitors leading to increased numbers of neutrophils and monocytes, likely driven by increased levels of G-CSF (Gordy et al., 2011). In contrast, the maintenance and longevity of neutrophils during embryogenesis are less well understood. Yet, published work suggests a different life span of fetal and adult neutrophils, with the E14.5 fetal liver harbouring only a few apoptotic cells (Liu et al., 2010) in comparison to the adult spleen analysed in *LysMC^re^; c-Flip^f/f^* mice and littermate controls (Gordy et al., 2011). Indeed, circulating neutrophils at E16.5 can be fate-mapped to an E8.5 yolk sac progenitor (Gomez Perdiguero et al., 2015), and during embryogenesis, there is rather a massive increase of neutrophils between E14.5 and E16.5 (Freyer et al., 2021) (and own data, not shown) instead of the steady-state turnover observed in adult mice. Thus, increased numbers of neutrophils in the fetal liver of *Tnfrsf11a^Cre/+^; Spi1^f/f^* embryos are unlikely caused by an increase of apoptotic neutrophils not being phagocytosed by macrophages.

Our RNA-seq data instead indicate that there is a transcriptional change in LT-HSCs when macrophages are lacking, supporting the hypothesis that fetal liver macrophages provide not only a niche for erythroblasts but also for LT-HSCs. In LT-HSC from *Tnfrsf11a^Cre/+^*; *Spi1^f/f^* fetal livers *Gata2* and *Gata3* were upregulated compared to littermate controls. Previous studies have shown the importance of *Gata2* and *Gata3* transcription factors in hematopoiesis (Alsayegh et al., 2019). GATA2 serves as a regulator of genes controlling the proliferative capacity of early haematopoietic cells during embryogenesis (Tsai et al., 1994; Tsai and Orkin, 1997) and the GMP cell fate (Rodrigues et al., 2008). Here, gene dosage is also crucial for HSC functionality since *Gata2* heterozygote (*Gata2^+/-^*) mice displayed reduced GMP numbers in the bone marrow and serial replating CFU assays of *Gata2^+/-^* bone marrow produced less granulocyte-macrophage progenitors compared to controls (Rodrigues et al., 2008). Further, a study in GATA2-deficient human embryonic stem cells could show that GATA2 is required for the production of granulocytes (Huang et al., 2015). These data highlight the conserved function of *Gata2* in regulating HSC functionality on different levels, including their differentiation into granulocytes. Similar to Gata2, Gata3 can also regulate HSC maintenance and differentiation. Different studies of fetal and adult HSCs demonstrated that Gata3 is required for maintaining the self-renewing capacity of HSCs (Fitch et al., 2012; Frelin et al., 2013; Ku et al., 2012) and that expression of Gata3 is tightly regulating LT-HSCs to either self-renew or differentiate (Frelin et al., 2013). We could not observe an impact of *Gata2/Gata3* upregulation on LT-HSC numbers or their proliferation/serial replating capacity, suggesting a different mechanism in fetal livers. However, dysregulation of *Gata2* and/or *Gata3* expression may cause increasing numbers of GMPs and significantly more neutrophils in the fetal liver observed in *Tnfrsf11a^Cre/+^; Spi1^f/f^* embryos. To define the LT-HSC niche in more detail, it will be important to examine whether macrophage-derived signals can control the expression and/or activity of Gata2 and Gata3 or other transcription factors that are known to control HSCs stem-cell ness and differentiation.

A study in zebrafish supports the notion of macrophage-HSC crosstalk requirement during development, uncovering a ‘grooming’ mechanism of embryonic macrophages that had a long-lasting impact on adult stem cells: HSPCs in the CHT often completed a cell division shortly after macrophage interactions and lack of CHT macrophages led to a decreased HSPC clonality in the adult marrow (Wattrus et al., 2022). Interestingly, their data suggests that HSPC proliferation in the CHT is mediated through ERK/MAPK activity, which is controlled by macrophage-derived Il1b (Wattrus et al., 2022). Indeed, the scRNA-seq macrophage cluster 2 is specifically expressing Il1b and could have a similar effect on a subset of LT-HSCs in the mouse fetal liver. Therefore, dissecting the macrophage-derived ligands and their effect on HSPC populations on a single-cell level will shed light on the crosstalk mechanisms in mouse fetal livers in the future.

Defining factors that control the HSC niche is essential to support efforts for an *in vitro* expansion and targeted differentiation of HSCs. Only a few studies have focused on macrophages as HSC niche cells so far, as they are primarily viewed as interacting cells of erythroblasts in both the fetal liver and the adult bone marrow. Here, we show that macrophages provide a niche for LT-HSCs in the fetal liver, and that macrophage deficiency leads to changes in the LT-HSC transcriptional program and their differentiation capacity. Likely, not only the immediate interaction but also macrophage-derived cytokines and growth factors affect processes, such as fate decisions and stem-cell ness. Our results provide a starting point to study the impact of macrophage-derived signals on LT-HSC functionality during embryogenesis and adulthood.

## Supplementary figure legends

**Supplementary figure 1.**
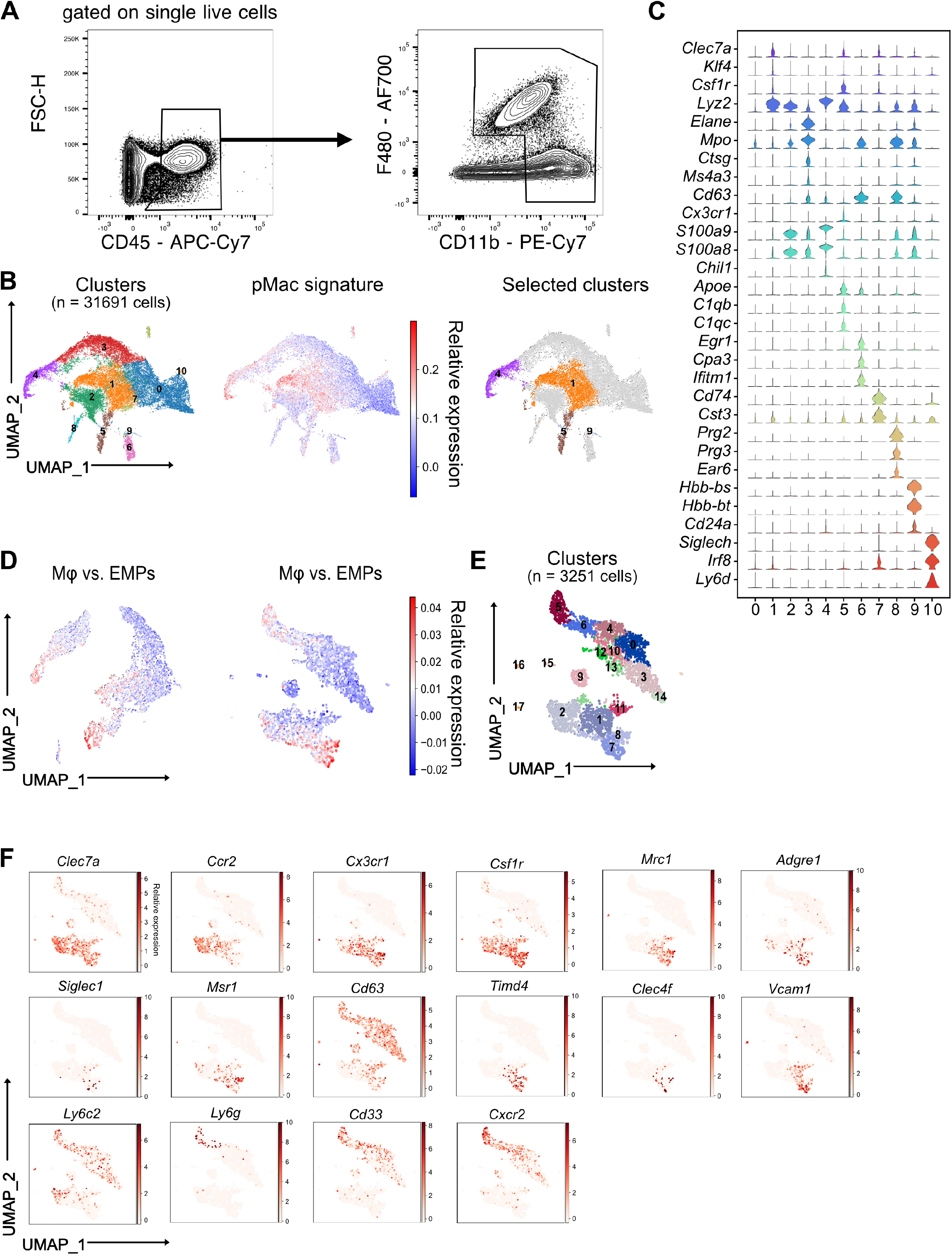
Sorting and characterization of fetal liver myeloid cells. **(A)** Sorting strategy of CD11b^+^ cell for performing single-cell RNA-sequencing using wildtype E14.5 fetal livers. **(B)** Sub-clustering strategy of the single-cell clusters from all cells passing quality control. The initial clusters of interest were identified by projecting the pre-macrophage (pMac) signature on the single cells. **(C)** Top expressed genes from clusters in (B) to identify myeloid cell populations. **(D)** Sub-clustering strategy for the second and the third round of macrophage cluster selection. The initially selected clusters were analysed for a second time, and a set of signature genes upregulated in macrophages (Mφ) compared to erythro-myeloid progenitors (EMPs) were projected on them. The same procedure was repeated for a third time. **(E)** Clusters resulting from the third round of sub-setting. 18 identified clusters were visualized using UMAP. **(F)** Expression of individual monocyte and macrophage-specific genes on the final identified clusters.

**Supplementary figure 2.**
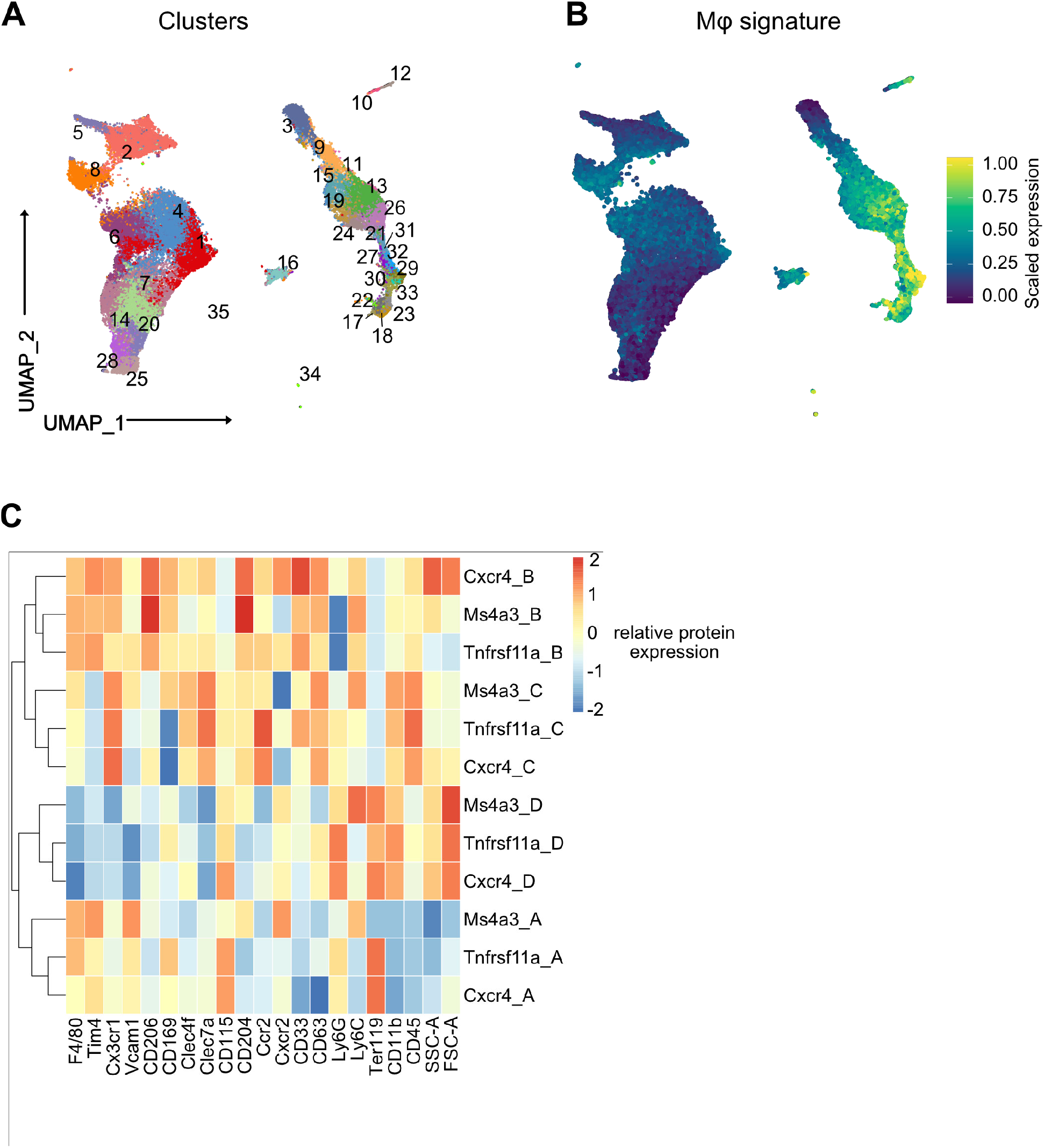
Flow cytometry analysis of macrophage subpopulations and fate-mapping mouse models. **(A)** Flow cytometry analysis of CD11b^low/+^ cells with a macrophage signature, isolated from a fetal liver at developmental day E14.5. Cell surface marker expression was used to generate unbiased clusters using UMAP. **(B)** Projection of macrophage markers on the flow cytometry data using UMAP visualization. **(C)** Relative expression of markers used in the flow cytometry analysis of three different fate-mapping mouse models. Tnfrsf11a: *Tnfrsf11a^Cre^; Rosa26^YFP^* model. Ms4a3: *Ms4a3C^re^; Rosa26^YFP^;* Cxcr4: *Cxcr4^CreERT^; Rosa26^YFP^* with 4-hydroxytamoxifen (4-OHT) injection at E10.5. The identified macrophage clusters (A-D) shown in Figure 1F could be identified in all models with similar expression patterns.

**Supplementary figure 3.**
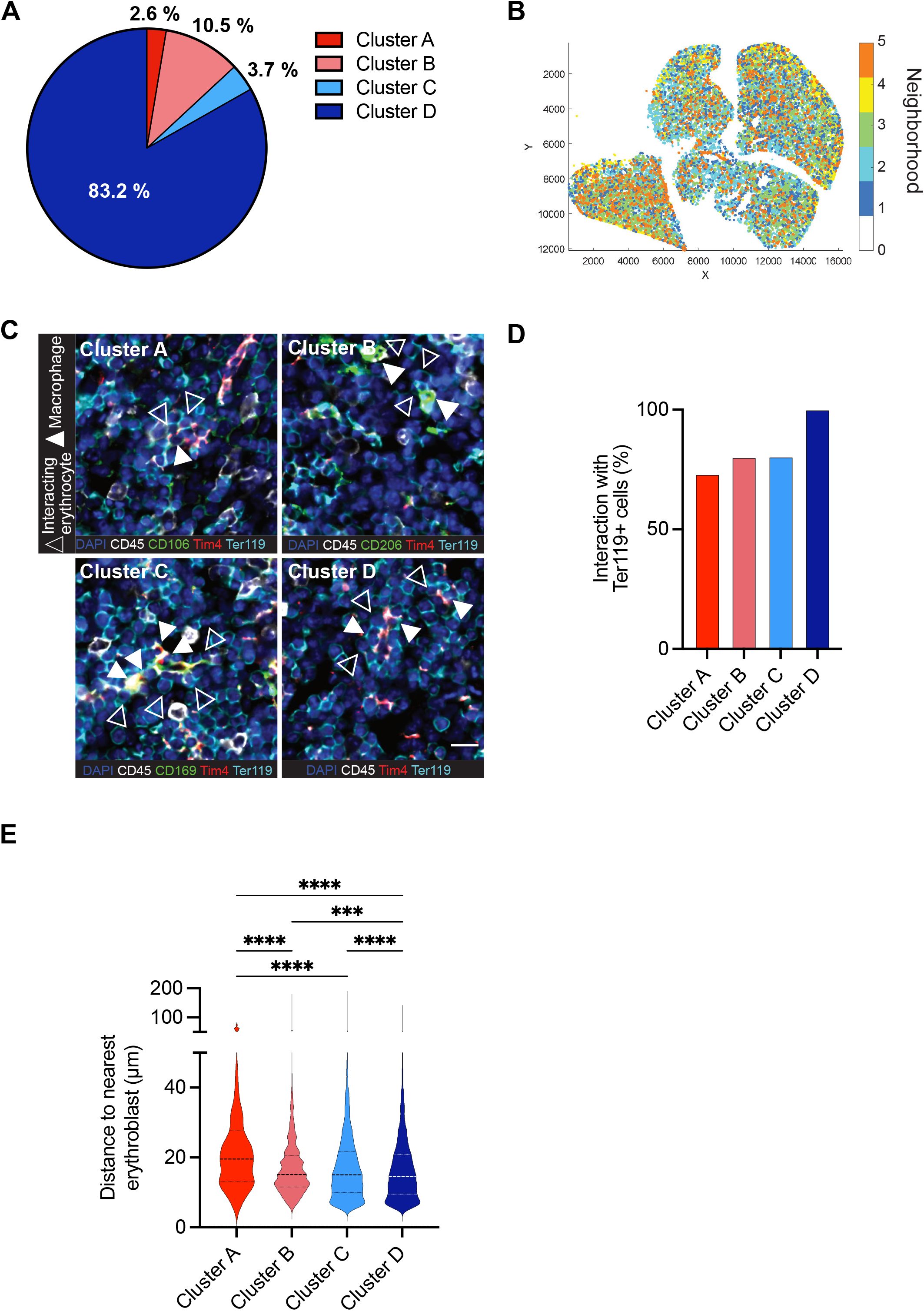
Spatial CODEX analyses of macrophage clusters in the fetal liver. **(A)** Proportions of cells from the macrophage clusters within the total F4/80^+^ Iba1 ^+^ Cx3cr1^+^ Tim4^+^ macrophages from the CODEX image in Figure 3B. **(B)** Single objects detected in Figure 3B were spatially segmented into neighbourhoods using a raster scan with a radius of 50μm. Plot represents clustering of cellular neighbourhoods based on their local composition of macrophages from the four clusters. Each colour represents a neighbourhood with a similar composition. **(C)** Representative images of interactions between macrophage subpopulations and Ter119^+^ erythroblasts. Filled arrowheads indicate macrophages from the corresponding cluster, whereas empty arrowheads indicate interacting erythrocytes. Scale bar represents 15 μm. **(D)** Interaction frequency between Ter119^+^ cells and distinct macrophage clusters, as indicated in (C). **(E)** The distance from the four macrophage clusters to their nearest erythrocyte in the entire tissue section was measured. One-way ANOVA; *** *p* < 0.001, **** *p* = 0.0001.

**Supplementary figure 4.**
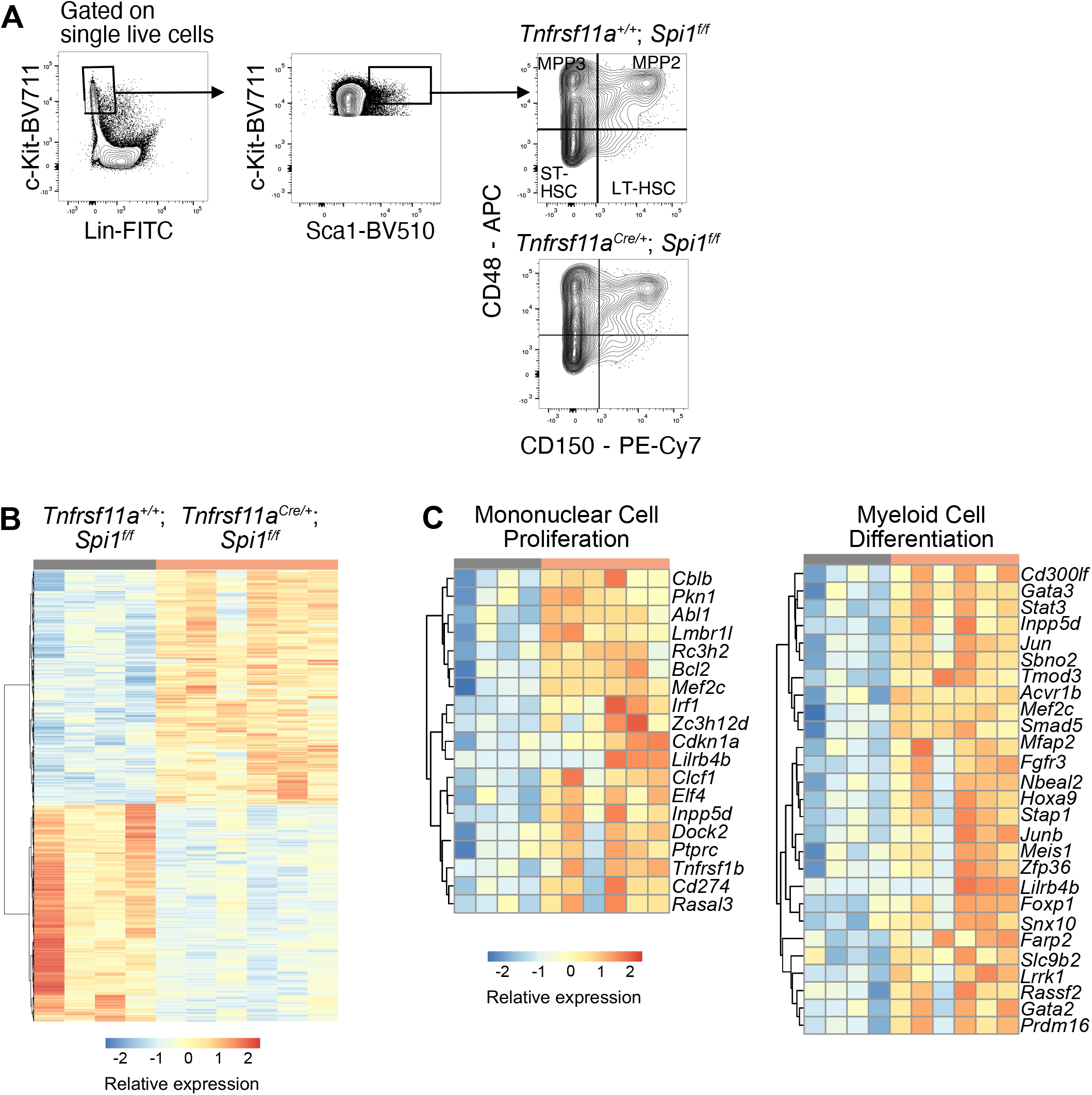
Bulk RNA-sequencing and analysis of LT-HSC. **(A)** Gating strategy for sorting LT-HSC from *Tnfrsf11a^+/+^; Spi1^f/f^* and *Tnfrsf11a^Cre^; Spi1^f/f^* fetal livers at E14.5 for RNA-sequencing. **(B)** Heatmap of all differentially expressed genes between LT-HSCs from *Tnfrsf11a^+/+^; Spi1^f/f^* and *Tnfrsf11a^Cre^; Spi1^f/f^* at E14.5 with adjusted p-value <0.1. **(C)** Heatmap of genes belonging to the GO terms ‘mononuclear cell proliferation’ and ‘myeloid cell differentiation’.

**Supplementary figure 5.**
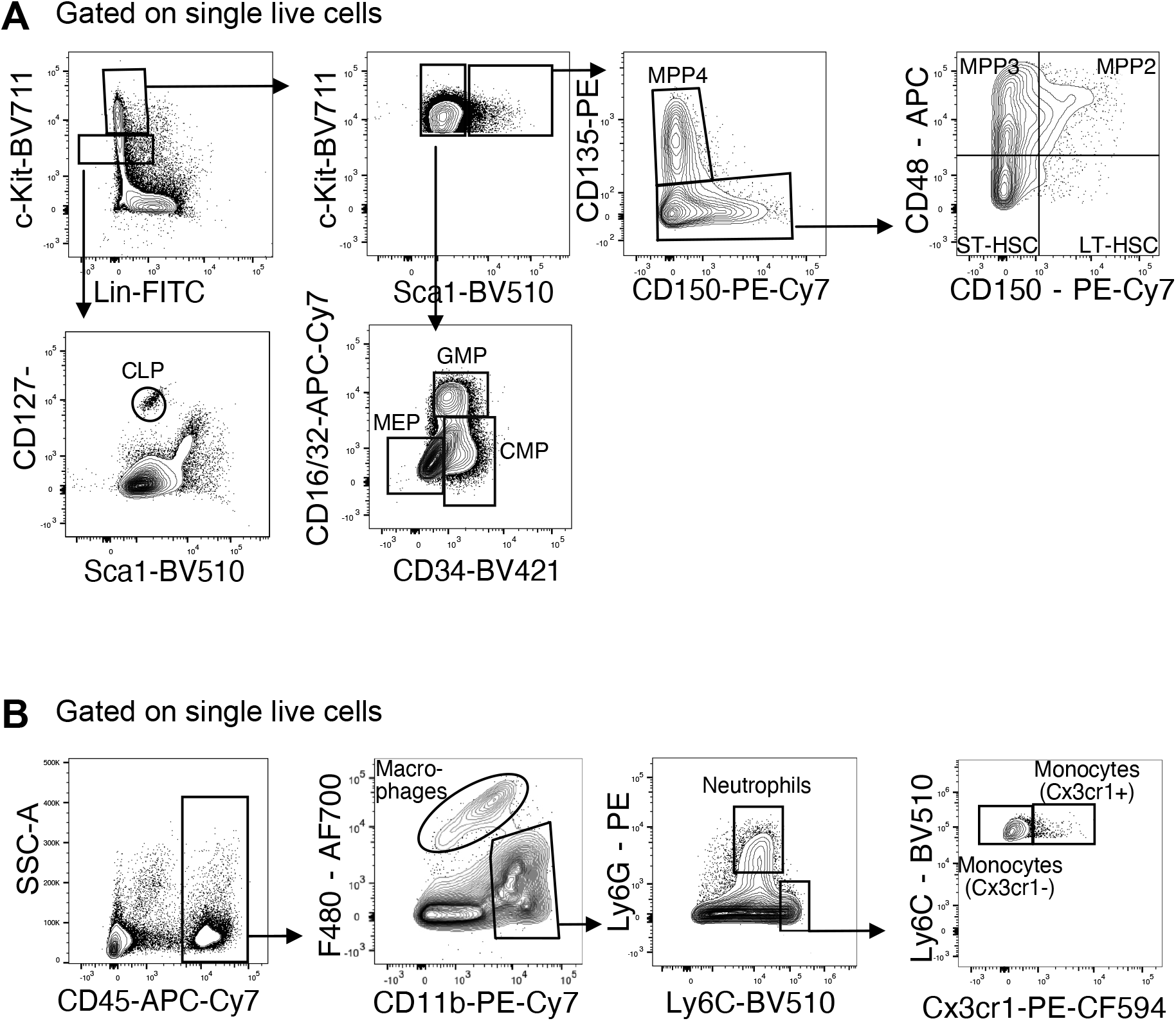
Gating strategies for flow-cytometry data. **(A)** Gating strategy for quantification of stem and progenitor cells in fetal livers at E14.5. CLP: common lymphoid progenitor; CMP: common myeloid progenitor; GMP: granulocyte-macrophage progenitor; LT-HSC: long-term hematopoietic stem cells; MEP: megakaryocyte-erythrocyte progenitor; MPP: multipotent progenitors; ST-HSC: short-term hematopoietic stem cells. **(B)** Gating strategy for quantification of myeloid cells in fetal livers at E14.5.

## Acknowledgements

We thank Cornelia Cygon for technical support and Florent Ginhoux for providing *Ms4a3^Cre^* mice. The work in the labs was supported by the following grants: a) Funded by the Deutsche Forschungsgemeinschaft (DFG, German Research Foundation) under Germany’s Excellence Strategy-EXC2151-390873048 (to EM, JLS, EK, and AS), GRK2168 (to EM), GRK1873/2 (to EM), SFB1454 (to EM, JLS, AS, and MB), b) Boehringer Ingelheim Fonds (doctoral fellowship to KM).

## Author contributions

EM and AHK conceived the project. AHK, IS, DAB, HH, KM, NM, DH, NRB, COS, performed experiments. AHK, IS, DAB, HH, DH analyzed data. EM, SU, MB, AS supervised experiments and data analysis. KB and JLS provided help with scRNA-seq experiments and analyses. EM, EK, SU, AS gave technical support and conceptual advice. EM and AHK wrote the manuscript. All authors discussed the results and commented on the manuscript.

## Material and Methods

### Chemicals and solutions

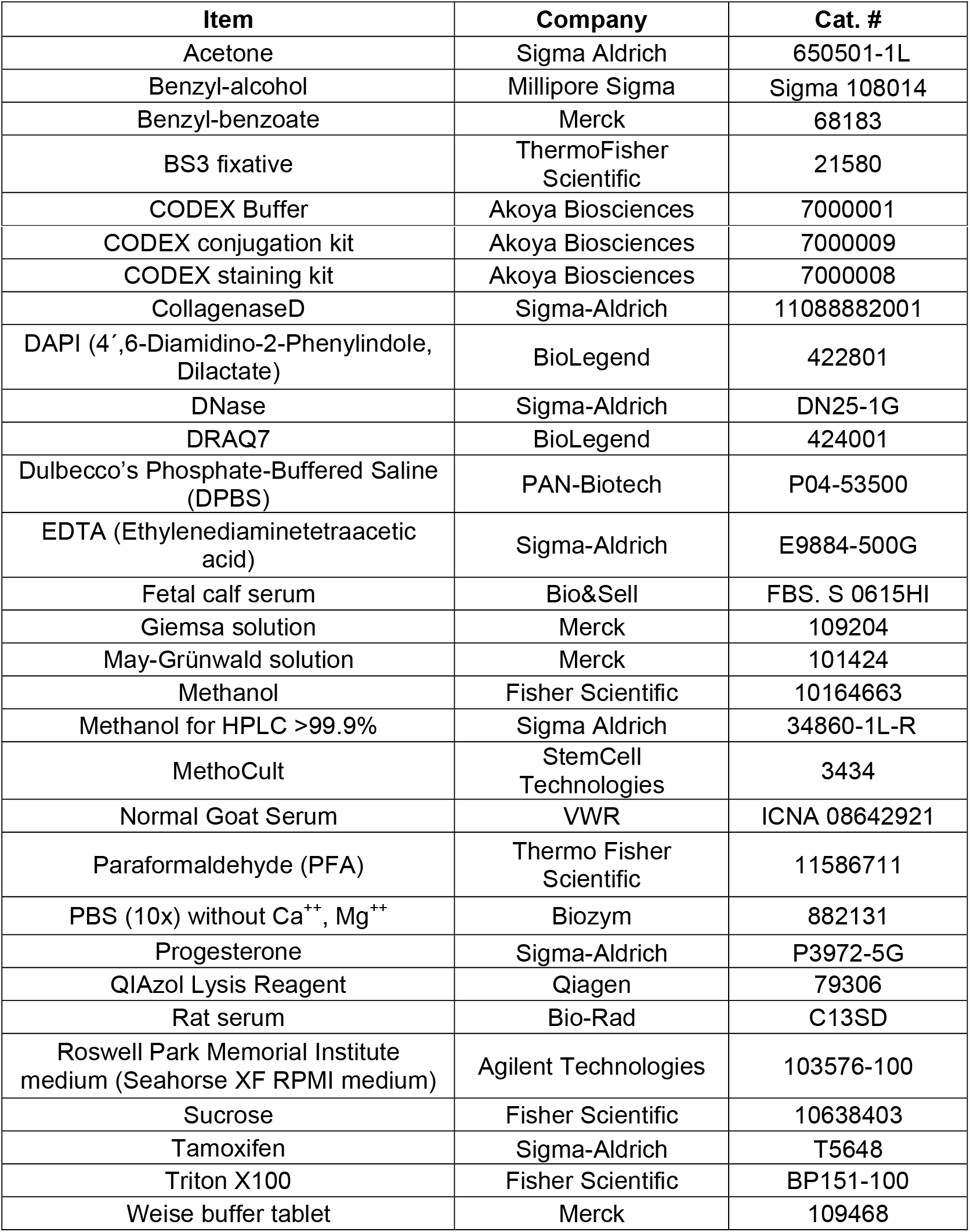

### Experimental mice

All mice were maintained on a C57BL/6 background and housed in SPF conditions. Animal procedures were performed in adherence to our project license 2018.A056 issued by the “Landesamt für Natur, Umwelt und Verbraucherschutz” (LANUV). Whenever possible, The ARRIVE guidelines 2.0 were followed (The ARRIVE Essential 10, https://arriveguidelines.org/arrive-guidelines). *Tnfrsf11a^Cre/+^; Spi1^f/+^* males were crossed to *Spi1^f/f^* females to generate embryos lacking macrophages. *Tnfrsf11a^Cre/+^, Ms4a3^Cre/+^*, and *Cxcr4^CreERT/+^* mice were crossed to the *Rosa26^YFP^* strain to generate embryos suitable for the fate-mapping. Adult mice were mated overnight to obtain embryos. To fate-map HSCs, *Cxcr4^CreERT^; Rosa26^YFP^* embryos were pulsed using 4-hydroxytamoxifen injection (75mg/kg) at embryonic day (E)10.5. To prevent tamoxifen-related abortions, progesterone (37.5mg/kg) was injected simultaneously with tamoxifen into the mice. The female was examined for vaginal plug formation the next day and the embryos were considered to be E0.5.

### Cryosection and whole-mount immunostaining

Pregnant mice were sacrificed through cervical dislocation. Embryos at E14.5 were harvested and stored in cold 1x Dulbecco’s Phosphate-Buffered Saline (DPBS w/o Ca and Mg, PAN-Biotech) and dissected under a Leica M80 microscope. Fetal livers (FL) were harvested and fixed in 1 % paraformaldehyde (PFA) overnight at 4 °C for immunofluorescence staining. Fixed lobes were washed three times for 10 min with DPBS, incubated in 30 % sucrose. The fixed lobes were dehydrated using increasing methanol (Fisher Scientific) gradient diluted in DPBS. Samples were incubated with primary antibodies (AB) (Table S5) overnight at 4 °C in 0.4 % PBT (DPBS with 0.4 % TritonX x100). Afterward, the samples were washed three times for 10 min in washing buffer (DPBS with 0.2 % Triton x100, 3 % NaCl (Grüssing) at room temperature (RT). The same procedure was repeated using the secondary antibodies (Table S5) and, if needed, using the directly conjugated antibodies a third time. samples were cleared in benzyl-alcohol benzyl-benzoate (BABB 1:2 proportion). Fetal liver lobes were placed for 30 min in 50 % BABB, followed by incubation in 100 % BABB for 30 min to obtain transparent tissues. The samples were placed in a cavity slide (Brand) filled with BABB. A round cover glass was carefully placed on the tissue and sealed using nail polish. The samples were 3D visualized using LSM 880 Zeiss confocal microscope with a 63x (oil) objective.

### Co-detection by indexing (CODEX)

5μm slices of fetal liver from E14.5 wildtype embryos were prepared and used for CODEX staining following the manufacturer’s instructions. Briefly, sections were retrieved from the freezer, let dry on drierite beads, and fixed for 10 min in ice-cold acetone (Sigma Aldrich, St. Louis, MO, USA). After fixation, samples were rehydrated and photobleached twice as described in (Du, Lin et al. 2019). Following photobleaching, sections were blocked and stained with a 20-plex CODEX antibody panel (Table S5 and S6) overnight at 4 °C. After staining, samples were washed, fixed with ice-cold methanol, washed with 1x PBS, and fixed for 20 min with BS3 fixative (Sigma Aldrich, St. Louis, MO, USA). A final washing step with 1x PBS was performed.

A multicycle CODEX experiment was performed following the manufacturer’s instructions. Images were acquired with a Zeiss Axio Observer widefield fluorescence microscope using a 20x objective (NA 0.85) and z-spacing of 1.5μm. The 405, 488, 568, and 647 nm channels were used. After imaging, raw files were exported using the CODEX Instrument Manager (Akoya Biosciences, Marlborough, MA, USA) and processed with CODEX Processor v1.7 (Akoya Biosciences). Image processing included background subtraction using the DAPI signals of the first and last empty cycles of the acquisition, deconvolution, shading correction, and stitching. For cell segmentation, DAPI counterstain was used for object detection, whereas sodium-potassium ATPase antibody staining was used as a membrane marker for delineating the cell shape.

A manual cell classification was performed in CODEX MAV 1.5 (Akoya Biosciences). Annotation of the macrophage clusters was done using the same gating strategy as in flow cytometry, with the difference that F4/80^+^CD11b^+^ cells were not gated but F4/80^+^ Iba1^+^ cells. HSCs were gated as CD150^+^ c-Kit^+^ cells, erythrocytes as CD45^-^Ter119^+^ cells, and blood vessels as CD45^-^CD31^+^SMA^+^ cells. After cell classification, Voronoi diagrams were generated in CODEX MAV using the four macrophage clusters, blood vessels, and HSCs as seeds.

### Spatial analyses and determination of cellular neighborhoods with CODEX images

LogOddRatio analysis for spatial interactions was performed in CODEX MAV. For this, after cell classification, the four macrophage populations, HSCs, and blood vessels were selected. The selected minimum and maximum distances of interaction were 5 and 50 μm, respectively.

For cellular neighborhood analyses, the .csv files generated with CODEX MAV were exported to CytoMAP (Stoltzfus, Filipek et al. 2020) and the same cell classification was used to annotate the cells. A raster scan with a radius of 50μm was performed to spatially segment the image. To define the cellular neighborhoods based on local composition, a self-organizing map (SOM) clustering algorithm was used, considering only the macrophage populations. Heatmaps were generated to determine the cell composition of each neighborhood. To measure the distances between macrophage clusters and erythrocytes, images were exported to QuPath v0.3. and cells were detected using DAPI signals. Single object classifiers for each marker were trained, and these were used to generate composite classifiers to identify macrophage populations, erythrocytes, and HSCs as before. The distance between the cells of each macrophage cluster and their closest erythrocyte was measured and plotted.

To validate the proximity of macrophage clusters to HSCs, images were exported to QuPath v0.3, cells were segmented, and HSCs were identified, as described above. A circle with a fixed radius of 50μm was drawn and centered on 20 randomly selected HSCs. Next, the same composite classifiers to identify macrophage clusters were applied to the annotated circles, and the number of cells of each macrophage cluster within the defined radius was counted.

### Software

| Software | Producer | Identifier |
| --- | --- | --- |
| CODEX instrument manager | Akoya Biosciences |  |
| CODEX Processor | Akoya Biosciences |  |
| CODEX MAV | Akoya Biosciences |  |
| QuPath |  | (Bankhead et al., 2017) |
| Image J |  | (Schindelin et al., 2012) |
| FlowJo | BD |  |

### Colony-forming unit assay

CFU assays were performed according to the Mouse “Colony-Forming Unit” (CFU) Assays Using MethoCult™” GF M3434 protocol (StemCell Technologies) (STEMCELL). Briefly, a fetal liver lobe was collected in fluorescence cell sorting (FACS) buffer (1x DPBS with 2 % 100 mM Ethylenediaminetetraacetic acid (EDTA, Sigma-Aldrich), 0.5 % BSA) and digested using digestion solution (1 % DNase (Sigma-Aldrich), 1 % collagenase D (Sigma-Aldrich), 3 % fetal calf serum (FCS, Bio&Sell), 1x DPBS). The samples were mechanically disrupted and incubated for 30 min at 37 °C. Following the digestion, the whole volume was transferred in a FACS tube and centrifuged for 5 min at 400 g, 4 °C. The supernatant was discarded, and the pellet was resuspended in 1 ml sterile Roswell Park Memorial Institute medium (RPMI, supplemented with 10 % FCS, 1 % Penicillin-Streptomycin, 1 % D-Glutamate, 1 % Pyruvate). 3×10^5^ live cells were taken from the suspension and filled up to 1 ml with RPMI to achieve a 3×10^5^ cells/ml concentration. 1.5 ml MethoCult aliquots (StemCell Technologies) were thawed at room temperature and vortexed vigorously. The cell suspension was added to the MethoCult in a way to achieve 3×10^4^ cells/ml concentration. 1 ml MethoCult mixed with the cell suspension was slowly taken up using a pipette and then transferred to a 35 mm cell culture dish (VWR). The cell culture dish was cautiously tilted until it was covered with medium and was put inside the incubator. After 12 days of cell culture, colonies were first identified by their phenotype, i.e. their size, shape, and density were analyzed based on representative pictures shown in the CFU assay protocol by STEMCELL Technologies. Subsequently, colonies were picked and identification was validated by the May-Grünwald-Giemsa staining (Merck) of the colonies. Prior to the staining, the colonies were picked up using a 10 μL pipet under Leica M80 microscope and collected into microtubes containing 10 μL FACS buffer. The collected colonies were transferred on slides using cytospin funnels (Hettich), centrifuged at 800RPM by a cytospin centrifuge (Hettich) for 10 min. After the centrifuge the slides were air-dried and fixed using cold methanol.

### Flow-cytometry sample preparation and data acquisition

Pregnant mice were sacrificed through cervical dislocation at E14.5. The fetal liver, brain, and lung were harvested from the embryos of the *Tnfrsf11a^Cre^; Spi1^flox/+^*. The collected tissues were digested for 20 min at 37 °C in a digestion solution. The cell suspensions were centrifuged and the supernatants were removed. The pellets were resuspended in 50μL blocking solution (2 % rat serum) for 10 min incubation on ice. The volume of each sample of the *Tnfrsf11a^Cre^; Spi1^flox/+^* model was measured to obtain the total cell number. The samples were stained with primary antibodies for 25 min (Table S5). Afterward, the samples were washed by adding 100 μL FACS to the suspension and centrifuge at 400 g for 5 min at 4 °C. The same procedure was repeated for the secondary antibodies. Finally, the cells were stained with Hoechst live/dead staining (1:10000) before flow-cytometry and recorded using a FACSymphony™ (BD Biosciences) cytometer.

The same procedure was also done with harvested fetal livers from wild-type embryos at E14.5. These samples were stained with primary and secondary antibodies designated to investigate the heterogeneity of macrophages (Table S5). The cells were stained with DRAQ7 live/dead staining (1:1000) before flow-cytometry and recorded using a FACSymphony™ cytometer.

### Analysis of flow-cytometry data for quantification of cells

Flow-cytometry data analysis was performed using FlowJo™ Software v.10.8.1 Becton, Dickinson, and Company. the quantification of myeloid cells in the fetal liver and the erythrocytes differentiation was done using the supplementary Figure 3A gating strategy. The quantification of HSCs and progenitors was done using supplementary Figure 3B gating strategy and the quantification of macrophages in the control organ, the fetal brain, and the fetal lung, was done using supplementary Figure 3C gating strategy. The count of each cell type was recorded and plotted using R (v. 4.0.5) and the ggplot2 (v. 3.3.5) and ggpubr (v. 0.4.0) (Wickham, Chang et al. 2016, Kassambara and Kassambara 2020).

### Analysis of flow-cytometry data for heterogeneity of macrophages

The CD11b^+^ F4/80^+^ cells were gated (Figure S1A) and downsampled using downsample plug-in (v.3.3.1) in Flowjo. The downsampled population was imported and analyzed in R (https://www.r-project.org/ v.4.0.5). The importing and processing of data was done using the CATALYST package (v. 1.18.1) (Crowell H 2022), which was installed through the Bioconductor package (v 3.14). The visualization of data was done using the UMAP algorithm (McInnes, Healy et al. 2018), and the clustering of data was done using FlowSOM (Van Gassen, Callebaut et al. 2015) clustering and ConsensusClusterPlus metaclustering (Wilkerson and Hayes 2010). The resulting clusters were manually inspected for expression of different markers and clusters of interests were subset and merged if necessary to form final clusters that would represent the macrophages and their heterogeneity.

### Cell-sorting of HSCs and macrophages for RNA-sequencing

Pregnant mice were sacrificed through cervical dislocation at embryonic day E14.5. The fetal liver was collected from *Tnfrsf11a^Cre/+^; Spi1^f/f^* and wild-type mouse models’ embryos. The collected tissues were digested for 20 min at 37 °C in a digestion solution. The cell suspensions were centrifuged and the supernatants were removed. The pellets were resuspended in 50 μL blocking solution (2 % rat serum) for 10 min incubation on ice. The samples were washed and stained for 25 min. At the end of incubation, the samples were washed and centrifuged at 400 g for 5 min at 4 °C. Finally, the pellet was resuspended in FACS buffer. The cells were stained with DAPI live/dead staining in a final dilution 1:10000 before flow-cytometry analysis using BD FACS ARIA III^TM^. ~700-1200 LT-HSCs were sorted according to the gating strategy (Supplementary Figure 4A) into microtubes containing 500 μL of Qiazol lysis buffer while the CD11b^low/+^ F4/80^+^ cells (Supplementary Figure 1A) were sorted into microtubes containing 100 μL of FACS buffer. The cells were used for loading the arrays in the next steps for single-cell RNA sequencing.

### Bulk-RNA sequencing and analysis

10 samples in total (4 controls and 6 knockouts) were analysed for the bulk RNA-sequencing. Total RNA was extracted using the miRNeasy micro kit (Qiagen) and quantified via RNA assay on a tape station 4200 system (Agilent). 5ng total RNA was used as an input for library generation via SmartSeq 2 (SS2) RNA library production protocol as previously described (Picelli et al., 2014). Pre-amplification PCR was performed with 16 cycles for samples. Libraries were quantified using the Qubit HS dsDNA assay (Invitrogen), and fragment size distribution was analyzed via D1000 assay on a tape station 4200 system (Agilent). SS2 libraries were sequenced single-end with 75 cycles on a NextSeq2000 system using P3 chemistry (Illumina). Samples were demultiplexed and fastq files were generated using bcl2fastq2 v2.20 before alignment and quantification using Kallisto v0.44.0 based on the Gencode (mm10, GRCm38) vM16 (Ensembl 91) reference genomes. Sequencing results from the experiments were pseudoaligned using the Kallisto tool set. The counts were imported into R and analyzed using the DEseq2 package (Love, Huber et al. 2014). The genes with less than 11 counts in all samples were removed. The counts were transformed using the variance Stabilizing Transformation (VST) function of the DEseq2 pipeline. The knockout samples were compared to the control samples during the analysis, and genes were ranked on the differential expression (LFC threshold of 0.1, adjusted p-value (BH) <0.1). The ranked gene list was divided into down and up-regulated genes (Logfold2change >0.5 and <-0.5) and used for gene ontology analysis using the clusterProfiler package (Yu, Wang et al. 2012). The DEGs were visualized using volcano plot using Enhancedvolcano package (Blighe 2018).

### Single-cell RNA sequencing

Seq-Well arrays were prepared as described by Gierahn et al. (Gierahn, Wadsworth et al. 2017) . Seq-Well libraries were generated as described by Gierahn et al and Hughes et .al (Gierahn, Wadsworth et al. 2017, Hughes, Wadsworth et al. 2020). The cDNA libraries (1 ng) were tagmented with home-made single-loaded Tn5 transposase in TAPS-DMF buffer (50 mM TAPS-NaOH (pH 8.5), 25 mM MgCl_2_ 50 % DMF in H2O) for 10 min at 55 °C and the tagmented products were cleaned with the MinElute PCR kit following the manufacturer’s instructions. Finally, a master mix was prepared (2X NEBNext High Fidelity PCR Master Mix, 10 μM barcoded index primer, 10 μM P5-SMART-PCR primer) and added to the samples to attach the Illumina indices to the tagmented products in a PCR reaction (72 °C for 5 min, 98 °C for 30 s, 15 cycles of 98 °C for 10 s, 63 °C for 30 s, 72 °C for 1 min). The pools were cleaned with 0.6 x volumetric ratio AMPure XP beads. The final library quality was assessed using a High Sensitivity DNA5000 assay on a Tapestation 4200 (Agilent) and quantified using the Qubit high-sensitivity dsDNA assay. Seq-Well libraries were equimolarly pooled and clustered at 1.4pM concentration with 10% PhiX using High Output v2.5 chemistry on a NextSeq500 system. Sequencing was performed paired-end using custom Drop-Seq Read 1 primer for 21 cycles, 8 cycles for the i7 index, and 61 cycles for Read 2. Single-cell data were demultiplexed using bcl2fastq2 (v2.20). Fastq files from Seq-Well were loaded into a snakemake-based data pre-processing pipeline (version 0.31, available at https://github.com/Hoohm/dropSeqPipe) that relies on the Drop-seq tools provided by the McCarroll lab (Macosko, Basu et al. 2015). STAR alignment within the pipeline was performed using the murine GENCODE reference genome and transcriptome mm10 release vM16 (Team 2014).

### Analysis of single-cell RNA sequencing

The Single-cell RNA was analyzed using the scanpy package (v.1.8.1) (Wolf, Angerer et al. 2018) in python (v.3.4.1) (Van Rossum and Drake Jr 1995). The cells were pre-processed and filtered by checking for cells expressing less than 200 genes in less than three cells. After the filtration, cells were processed further and clustered using the Leiden algorithm (Traag, Waltman et al. 2019). The clusters were investigated using their differentially expressed genes (DEG). The DEG lists were identified using Wilcoxon-Signed-Rank Test (Rey and Neuhäuser 2011) by comparing each cluster to the rest of the clusters. The selection of the clusters of interest was done in an iterative way. Three clustering steps were performed in total to subset the cells. Briefly, the first round of clustering was done in a none-stringent manner and the resulted groups were monitored for their DEGs and were assigned to certain cell types that they resembled most. Based on the DEGs the clusters 1, 4, 5, and 9 were containing macrophages while the other groups were representing other cell types and states. This selection was assured further by exploring the expression of a set of pre-macrophages (pMac) markers among the selected clusters. Clusters that could contain macrophages were subset for a second round of clustering. The subset cells were processed and clustered for a second time to have more homogenous cells regarding cell types. Again, the clusters were subjected to another round of selection using two different sets of signature markers for genes that are expressed in macrophages. The selected clusters in the second round were subset and the cells were processed for a final round of clustering resulting in distinguishable clusters of macrophage cells and their precursors from the rest. The finals clusters were analyzed and the DEGs for the were extracted A psuedotime analysis using the Partition-based graph abstraction (PAGA) method (Wolf, Hamey et al. 2019) was later performed to analyze the trajectory of these groups.

### Correlation matrix

To make a correlation between the single-cell RNA-seq data and the flow-cytometry data, the clusters of interest from both of the datasets were extracted. The expression values of mutual markers between them were scaled and normalized for each of the two datasets. The correlation between the two datasets was calculated using spearman’s rank correlation coefficient and results were visualized using the pretty heatmap package (Kolde and Kolde 2015) in R.

### Ligand-Receptor analysis

CellTalk database information (Shao et al., 2021) was download and the ligands list were explored among the expressed genes of the five final macrophage clusters and the ligands that were expressed were selected (208 ligands). A GO analysis using ClusterProfiler package was done on the selected ligand’s genes. Terms that were associated with the hematopoiesis were selected and the genes belonging to these terms were extracted. Genes with a minimum of five counts (single-cell data) among the three final macrophage clusters were chosen (100 ligands). Their corresponding receptors were taken from the CellTalk database. The expression of those receptors was explored in the bulk RNA-seq data from control LT-HSCs and the receptors that had more than average of 40 counts were selected. The ligand and receptor interactions were visualized using a circular plot using circlize package (Gu, Gu et al. 2014).

### Data availability

RNA-seq data from bulk and single-cell experiments are available under GEO accession number GSE225444. Source data for CODEX pictures and analyses are available as pyramidal file at Drayd (Mass, Elvira (2023), Source Data Kayvanjoo et al., Dryad, Dataset, https://doi.org/10.5061/dryad.fn2z34v00). All other data will be made available upon reasonable request.

